# Inhibition of Parkinson’s Disease-related LRRK2 by type-I and type-II kinase inhibitors: activity and structures

**DOI:** 10.1101/2023.09.07.556689

**Authors:** Marta Sanz Murillo, Amalia Villagran Suarez, Verena Dederer, Deep Chatterjee, Jaime Alegrio Louro, Stefan Knapp, Sebastian Mathea, Andres E Leschziner

**Affiliations:** Department of Cellular and Molecular Medicine, School of Medicine, University of California, San Diego, La Jolla, CA 92093; Aligning Science Across Parkinson’s (ASAP); Institute of Pharmaceutical Chemistry, Goethe-Universität, Frankfurt 60438, Germany; Structural Genomics Consortium (SGC), Buchmann Institute for Life Sciences, Goethe-Universität, Frankfurt 60438, Germany; Department of Molecular Biology, School of Biological Sciences, University of California, San Diego, La Jolla, CA 92093

## Abstract

Mutations in Leucine Rich Repeat Kinase 2 (LRRK2) are a common cause of familial Parkinson’s Disease (PD), and a risk factor for the sporadic form. Increased kinase activity has been shown in both familial and sporadic PD patients, making LRRK2 kinase inhibitors a major focus of drug development efforts in PD. Although significant progress has been made in understanding the structural biology of LRRK2, there are no available structures of LRRK2 inhibitor complexes. To this end, we solved cryo-EM structures of LRRK2, wild-type and PD-linked mutants, bound to the LRRK2-specific type-I inhibitor MLi-2 and the broad-spectrum type-II inhibitor GZD-824. Our structures revealed LRRK2’s kinase in the active-like state, stabilized by type-I inhibitor interactions, and an inactive DYG-out type-II inhibitor complex. Our structural analysis also showed how inhibitor-induced conformational changes in LRRK2 are affected by its autoinhibitory N-terminal repeats. The structural models provide a template for the rational development of LRRK2 kinase inhibitors covering both canonical inhibitor binding modes.

## Introduction

Mutations in Leucine-rich repeat kinase 2 (LRRK2) are one of the most common drivers of the familial form of Parkinson’s Disease (PD) (*1–3*), and are also associated with increased risk for the sporadic form of PD (*1, 4*). The most frequent PD-linked mutations in LRRK2 increase its kinase activity (*5–8*), but hyperactivation of an otherwise wild-type LRRK2 has also been reported in idiopathic PD (*9*). As a result, LRRK2 has become a major target for the development of kinase inhibitors as therapeutics for PD (*10, 11*).

LRRK2 is a large, 2527-residue multi-domain protein. Its N-terminal half is comprised of armadillo (ARM), ankyrin (ANK) and leucine-rich (LRR) repeat domains, while its C-terminal half contains two catalytic domains: a GTPase (Ras Of Complex, or ROC), and a Ser/Thr kinase domain, as well as two architectural domains, a C-terminal Of ROC (COR) between the GTPase and the kinase, and a WD40 domain at the end (**Figure 1A**). Recent studies using cryo-EM have described structures for the C-terminal half of LRRK2 (“LRRK2^RCKW^”) (*12*), full-length LRRK2 alone (*13*) and bound to Rab29 (*14*), and both LRRK2 (*15*) and LRRK2^RCKW^ (*16*) bound to microtubules. Although a recent study proposed structures of LRRK2^RCKW^ bound to inhibitors generated using molecular dynamics (*17*), there are no published experimental structures of inhibitor-bound LRRK2, an important gap in our understanding of this protein as a therapeutic target.

**Figure 1.**
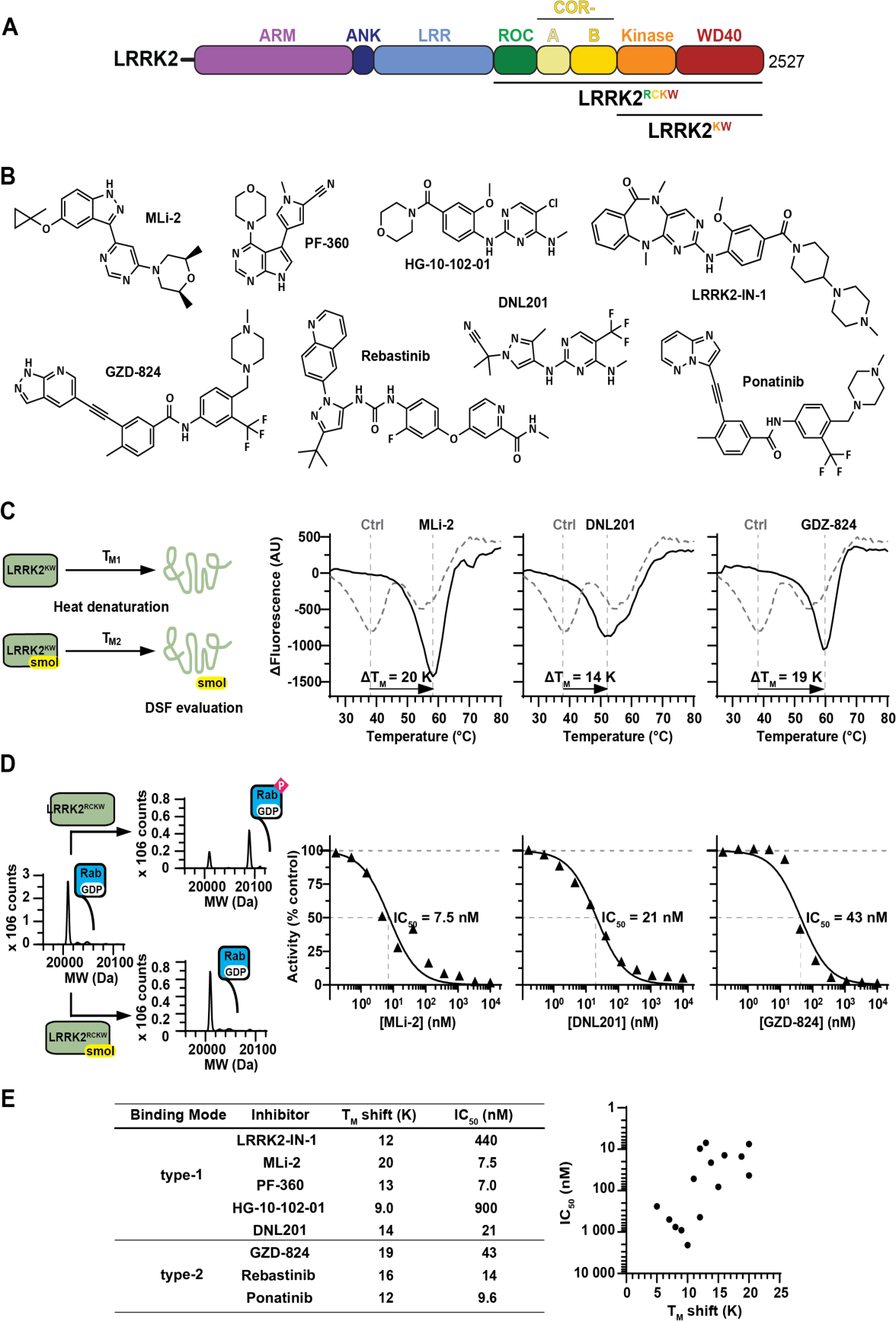
Stabilization and inhibition of LRRK2’s kinase by type-I and type-II inhibitors. **(A)** Domain architecture of LRRK2. The color coding of domains is used throughout this work. The three constructs used in this study—LRRK2, LRRK2^RCKW^, and LRRK2^KW^—are indicated. **(B)** Prominent inhibitors of the LRRK2 kinase. **(C)** The binding of kinase inhibitors stabilized LRRK2^KW^ as determined in a DSF assay. Shown are the first derivatives of the melting curves. The biphasic melting of LRRK2^KW^ indicated that the kinase and WD40 domains melt independently of each other. **(D)** The impact of type-I and type-II inhibitors on LRRK2^RCKW^ kinase activity was assessed with an MS-based activity assay. The inhibitors prevented LRRK2^RCKW^ from phosphorylating Rab8A with nanomolar IC_50_ values. **(E)** Correlation between T_M_ shift and IC_50_ for the inhibitors shown used in our assays (B).

### LRRK2 kinase inhibitors

Small molecules of diverse chemotypes have been developed as type-I kinase inhibitors for LRRK2 (**Figure 1B**). The first inhibitor to be introduced with an acceptable selectivity profile was LRRK2-IN-1 (*18*). However, its use for *in vivo* applications has been limited because of its low brain penetrance. HG-10-102-01, in contrast, was developed as a brain penetrant inhibitor (*19*), and its amino-pyrimidine based scaffold recently experienced a renaissance as a PROTAC warhead (*20*). MLi-2, an inhibitor with high affinity and a superb selectivity profile was a game changer (*21*). Ever since its introduction in 2015 it has been regarded as the gold standard LRRK2 inhibitor and this chemical probe has been used in numerous studies, including clinical trials. Another powerful but less prominent inhibitor is PFE-360 (*22*). Similar to MLi-2, its affinity for LRRK2 and its selectivity profile are both excellent. The last entry in our incomplete list is the type-I inhibitor DNL201 (developed as GNE0877) (*23*). This compound has been shown to efficiently inhibit LRRK2 in PD patients (*24*), and the related molecule DNL151 (renamed BIIB122) has currently entered phase 2b clinical testing (clinicaltrials.gov).

9/7/23 8:02:00 AMLess effort has been undertaken to develop type-II inhibitors for the LRRK2 kinase domain. To date, no selective type-II inhibitors have been published (*25*). However, several type-II inhibitors with a broad kinase target spectrum have been shown to bind to LRRK2 kinase with high affinity, namely ponatinib, GZD-824 (*12*) and rebastinib (*26*). However, their numerous kinase off-targets prevent their application as chemical tools or for the treatment of PD.

The need for LRRK2 selective type-II inhibitors is not limited to their therapeutic potential. The recent structural studies of LRRK2 listed above (*12–16*) have made it clear that conformational changes in LRRK2 driven by its kinase are likely to play central roles in its activation and subcellular localization. Having reagents that stabilize the LRRK2 kinase domain in either a closed, active-like conformation (type-I inhibitors), or in its open, inactive conformation (type-II inhibitors) would have a major impact on our ability to dissect its catalytic and scaffolding functions in cells and *in vivo*.

Here, we set out to bridge the gap in our understanding of the structural biology of LRRK2 inhibition. We present cryo-EM structures of LRRK2^RCKW^, WT and carrying the PD-linked mutations G2019S and I2020T, bound to the LRRK2-specific type-I inhibitor MLi-2 or the broad-spectrum type-II inhibitor GZD-824. We also present structures of full-length LRRK2 bound to these inhibitors to understand how the presence of the N-terminal repeats in LRRK2 affect inhibitor-induced conformational changes in the context of the full-length protein. The work presented here should help in the design of the next generation of LRRK2-specific inhibitors.

## Results

### Inhibitor binding stabilizes the LRRK2 kinase domain

The tight binding of small molecules to a protein usually stabilizes the protein and increases the temperature at which it denatures. We characterized the binding of diverse type-I and type-II inhibitors (**Figure 1B**) by measuring the thermostability increase of a truncated LRRK2 protein consisting of its kinase and WD40 domains (“LRRK2^KW^”) (**Figure 1A**) in the presence of inhibitors using differential scanning fluorimetry (DSF). Interestingly, the melting profile showed two clearly separated transitions most likely corresponding to the two domains, the kinase and the WD domain. In agreement with this hypothesis, only the melting curve of one of the phase transitions shifted in the presence of the diverse inhibitors tested suggesting that the transition at lower melting temperature corresponds to the kinase domain. However, in the absence of inhibitors, LRRK2^KW^ was relatively unstable compared to other kinases (melting temperature (T_M_) of 39°C) (*27*). Both type-I and type-II inhibitors significantly stabilized LRRK2^KW^ with T_M_ shifts of up to 20°C, indicating strong binding to the kinase domain, while the melting curve of the WD domain remained unaffected (**Figure 1C, E**).

### A mass spectrometry-based assay to monitor LRRK2 activity and inhibition

Increased kinase catalytic activity or impaired autoinhibition are features of LRRK2 PD variants (*5–8*). To compare the activity of the selected LRRK2 inhibitors, we developed a mass spectrometry (MS)-based kinase activity assay using an established endogenous substrate. We subjected the Rab8A GTPase domain (residues 6 to 176) to LRRK2^RCKW^ phosphorylation in the presence of varying concentrations of a given inhibitor and directly quantified the amount of phosphorylated Rab8A (pRab8A) by ESI-MS (**Figure 1D**). Most type-I and type-II inhibitors inhibited the phosphorylation reaction with IC_50_ values between 7 and 50 nM. Surprisingly, the inhibitors HG-10-102-01 and LRRK2-IN-1 stood out with IC_50_ values in the high nanomolar range. Altogether, there was a good correlation between the thermal shift data and the IC_50_ values (**Figure 1E**).

### Cryo-EM structures of LRRK2^RCKW^(G2019S) bound to type-I and type-II inhibitors

To understand how inhibitors of the two main canonical types interact with the catalytic domain of LRRK2, we began by determining cryo-EM structures of LRRK2^RCKW^, a well characterized construct consisting of the C-terminal half of LRRK2 containing all catalytic domains (**Figure 1A**) (*12, 16, 17, 26*), bound to the LRRK2-specific inhibitor MLi-2 (type I) or the broad-spectrum GZD-824 (type II), in the presence of GDP as a ligand for the GTPase (ROC) domain. We used a variant of LRRK2^RCKW^ carrying the G2019S mutation, the most common mutation associated with PD (*28*) and one of two, along with I2020T, located in the kinase domain (**Figure 2, Figures S2, S3, and S4, Table S1**).

**Figure 2.**
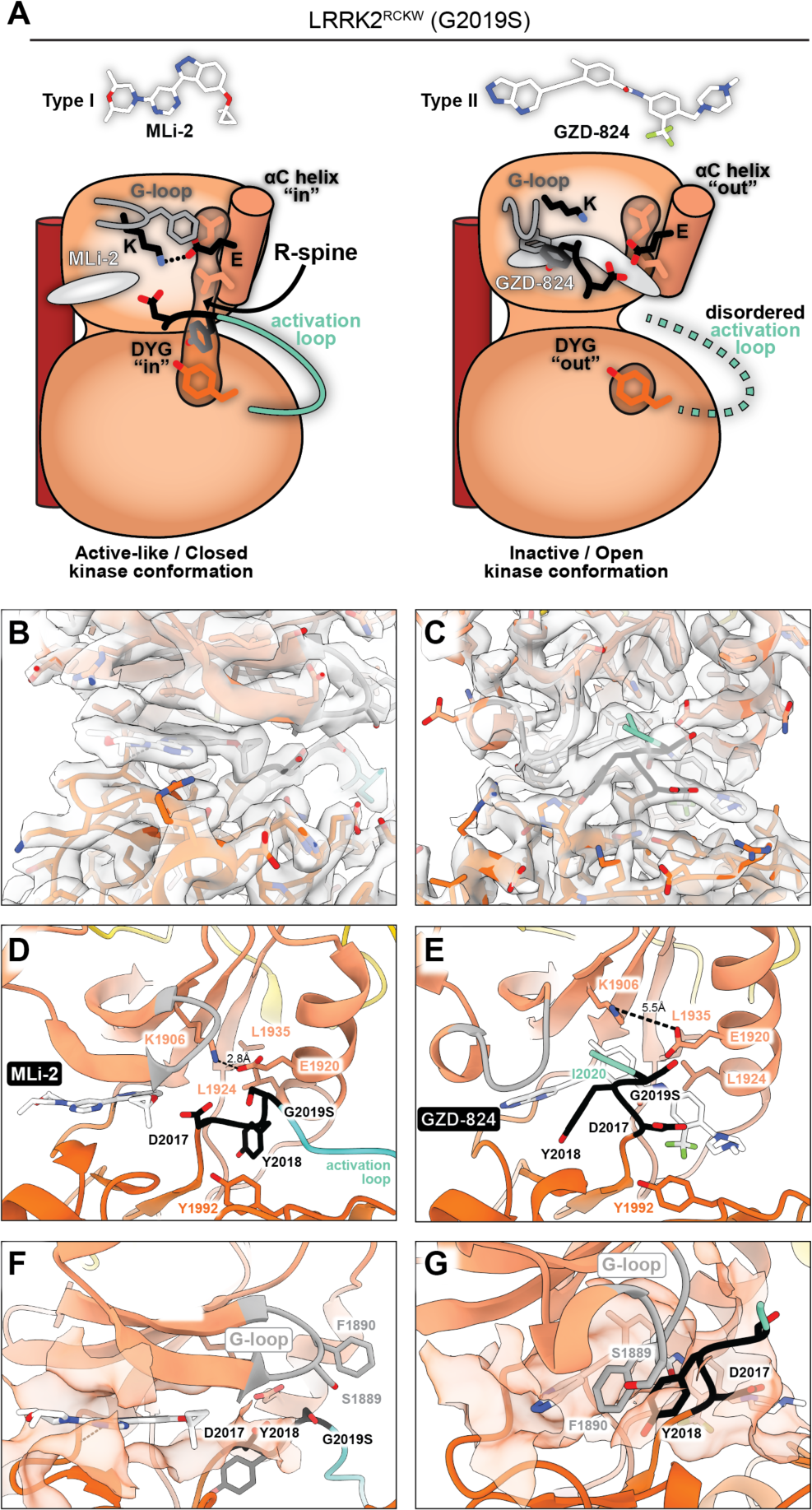
LRRK2^RCKW^(G2019S) bound to MLi-2 and GZD-824. **(A)** Cartoon summary of the main features of the structures presented here. The inhibitors used, the type-I MLi-2 and type-II GZD-824, are shown in stick representation. All panels below correspond to the structure depicted in cartoon form here. **(B,C)** Cryo-EM maps and models for the inhibitor binding site and surrounding regions for MLi-2 (B) and GZD-824 (C) bound to LRRK2^RCKW^(G2019S). **(D,E)** Close-up of the inhibitor binding site and kinase active site for the same structures shown in (B,C). Major features (G-loop, Lys1906-Glu1920 interaction, DYG motif, R-spine, and activation loop) are indicated. **(F,G)** Inhibitor binding pocket. The semitransparent orange surface indicates the solvent accessible surface for those residues in contact with the inhibitor. Some of the features highlighted in (D,E) are shown here as well.

We solved these structures, as well as all additional cryo-EM structures reported here, bound to a LRRK2-specific Designed Ankyrin Repeat Protein (DARPin) (*29*) that we refer to as “E11”. This DARPin was developed screening LRRK2^RCKW^, and it tightly bound to its WD40 domain (*manuscript in preparation*). We have found that adding the E11 DARPin to our cryo-EM samples consistently leads to structures with better resolution, most likely due to a reduction in the preferred orientation that LRRK2 and LRRK2^RCKW^ tend to adopt on cryo-EM grids, although we cannot rule out a contribution from their additional mass (*manuscript in preparation*). For simplicity, we will omit “E11” from the names of the complexes discussed in this work.

**Figure 2A** and **Movie S1** summarize the main features of the two structures. MLi-2 bound as expected in the ATP-binding pocket of LRRK2’s kinase, with its indazole group making hydrogen bonds with the backbones of E1948 and A1950 in the kinase hinge region, as recently proposed based on molecular dynamics simulations (*17*). In contrast to those simulations, however, the glycine rich (G) loop was extended in our structure, as expected for an active kinase domain and F1890, located at the tip of the G-loop, was not involved in coordinating MLi-2. Also as expected, MLi-2 binding stabilized the DYG “in” and αC “in” conformation, a fully formed R-Spine, and an ordered activation loop, (**Figure 2A,B**,D). (Note: even though we will continue using the “DYG” nomenclature throughout the paper, DYG is DYS in the G2019S mutant.) We observed an inactive conformation with DYG “out”, αC “out”, a broken R-Spine, and a disordered activation loop in the presence of GZD-824 (**Figure 2A,C,E**). Binding of GZD-824 to the ATP-binding pocket was shifted away from the hinge region relative to MLi-2; its pyrazolopyridine group partially overlapped with the location of the pyrimidine and indazole groups of MLi-2, with the rest of the molecule extending towards and past the αC helix. As expected from a larger inhibitor, the binding pocket for GZD-824 was enlarged by the DYG-out movement and opening of the allosteric back pocket, involving all the main components of the kinase’s active site (**Figure 2G**). A major difference between the two structures was the conformation of their G-loops (**Figure 2F,G**). While the Gloop was in an extended conformation in LRRK2^RCKW^(G2019S):MLi-2 (**Figure 2F**), it was sharply bent in LRRK2^RCKW^(G2019S):GZD-824, with F1890 interacting with the pyrazolopyridine group in GZD-824 (**Figure 2G**). In turn, Y2018 from the DYG motif interacted with F1890 (**Figure 2G**).

### Structural basis for decreased affinity of MLi-2 for LRRK2(G2019S)

G2019S is the most frequently identified LRRK2 mutation in PD patients (*28*). Although the G2019S mutation increases LRRK2 kinase activity, as do the other most common mutations (*5–8*), LRRK2(G2019S) is unusual: it is the only variant that does not increase microtubule association in cells (*30, 31*), it shows the highest levels of autophosphorylation at S1292 (*32*), and this mutation has been reported both to provide resistance to inhibition by MLi-2 (*33*), and to be more sensitive to this inhibitor (*25*). We therefore aimed to visualize, structurally, any differences that may exist in how inhibitors interact with LRRK2 carrying different PD-linked mutations. For this, we solved cryo-EM structures of LRRK2^RCKW^ wild-type (WT), and I2020T, bound to MLi-2 (**Figure 3, Figures S4, S5, and S6, Table S1**) and compared them to the structure of LRRK2^RCKW^(G2019S):MLi-2 presented above. We chose the I2020T mutation because of the different properties exhibited by G2019S and I2020T despite affecting neighboring residues.

**Figure 3.**
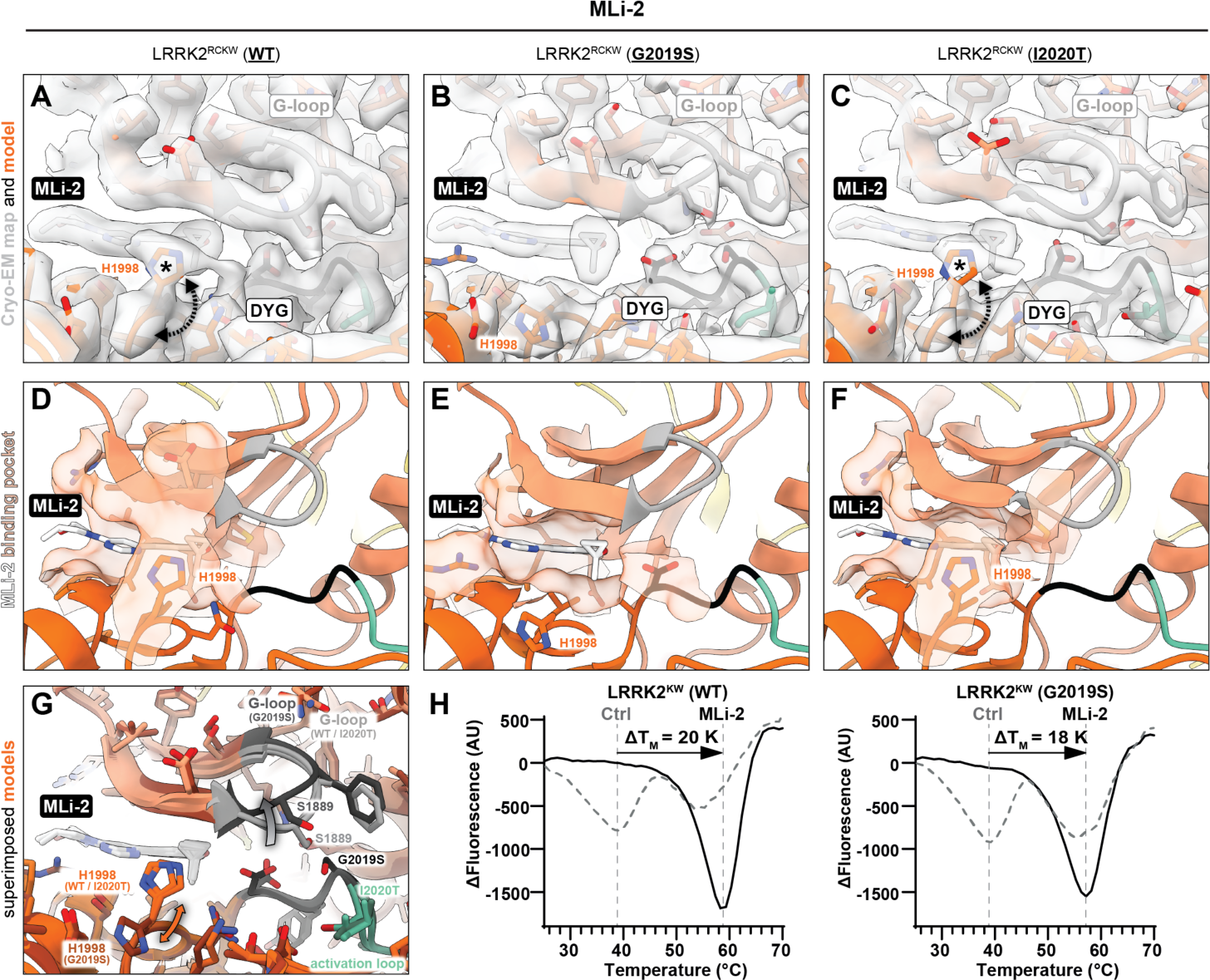
Binding of MLi-2 to LRRK2^RCKW^ wild-type (WT), G2019S, and I2020T. **(A-C)** Cryo-EM maps and models for the MLi-2 binding site and surrounding regions for LRRK2^RCKW^ WT (A), G2019S (B), and I2020T (C). The asterisks in (A) and (C) highlight densities present in the WT and I2020T maps but absent from the G2019S one. The dashed arrows in (A) and (C) indicate that two different rotamers of H1998 can be accommodated by the maps of WT and I2020T. The rotamer that points “up” (towards the N-lobe of the kinase) is shown in these panels. Note that only density accounting for the “down” rotamer is seen in G2019S (B). The G-loop and DYG motif are indicated in all three panels. **(D-F)** Inhibitor binding pocket. The semitransparent orange surface indicates the solvent accessible surface for those residues in contact with MLi-2. In the case of LRRK2^RCKW^ WT and I2020T, these surfaces were generated using the “up” rotamer of H1998. Note that the “up” rotamer of H1998 leads to a closing of the binding pocket in WT and I2020T (D, F) that is absent in G2019S (E). **(G)** Superposition of the models for MLi-2 bound to LRRK2^RCKW^ WT, G2019S, and I2020T. G2019S is shown in darker shades. The grey gradient-colored arrow in the G-loop indicates the movement of the loop in G2019S relative to WT and I2020T. The two-headed orange arrow indicates that two rotamers of H1998 in LRRK2^RCKW^ WT and I2020T can account for the density in our cryo-EM maps. **(H)** Differential Scanning Fluorimetry data measured for LRRK2^RCKW^(WT) and LRRK2^RCKW^(G2019S) in the presence and absence of MLi-2. In the absence of inhibitor, both proteins showed identical melting points suggesting that the mutant did not affect stability of the recombinant protein. Binding of the inhibitor resulted in a lower DT_M_ for LRRK2^RCKW^(G2019S), suggesting weaker binding affinity of MLi-2.

Although the three structures— WT, G2019S, and I2020T—were very similar (**Figure 3**), two features were unique to LRRK2^RCKW^(G2019S):MLi-2. First, the presence of a Ser instead of a Gly at position 2019 introduced a clash with the G-loop that pushed S1889 away from S2019 (**Figure 3A-C**,F). The second was an unexplained density we observed in the LRRK2^RCKW^(WT):MLi-2 and LRRK2^RCKW^(I2020T):MLi-2 cryo-EM maps, but not in that of LRRK2^RCKW^(G2019S):MLi-2 (**Figure 3A-C**). This additional density in WT and I2020T could accommodate an alternative rotamer for H1998 (**Figure 3A,B**,G), which closes the binding site around MLi-2 (**Figure 3D-F**). Given the absence of density for this rotamer in G2019S, we predicted weaker binding of MLi-2 to LRRK2’s kinase carrying the G2019S mutation. Although the accurate determination of sub-nanomolar inhibitor affinities for complex proteins is challenging, our DSF assay showed that the addition of MLi-2 to LRRK2^KW^(G2019S) resulted in less thermal stabilization than that observed with LRRK2^KW^(WT) (**Figure 3G**).

### Structures of LRRK2^RCKW^ bound to the type-II inhibitor GZD-824

Next, we solved cryo-EM structures of LRRK2^RCKW^ WT, G2019S, and I2020T bound to the broad-spectrum type-II inhibitor GZD-824 (**Figure 4, Figures S4, S7, and S8, Table S1**). As with MLi-2, the three structures—WT, G2019S, and I2020T—were very similar (**Figure 4G**). Although all three structures assumed a DYG “out”, αC “out” conformation with a broken R-Spine and a disordered activation loop, the WT cryo-EM map showed a closer distance between K1906 and E1920 (**Figure 4A,G**). In all three cases, the G-loop was folded over the inhibitor to interact with its pyrazolopyridine and Y2018 from the DYG motif, as described above (**Figure 4A-C**,G).

**Figure 4.**
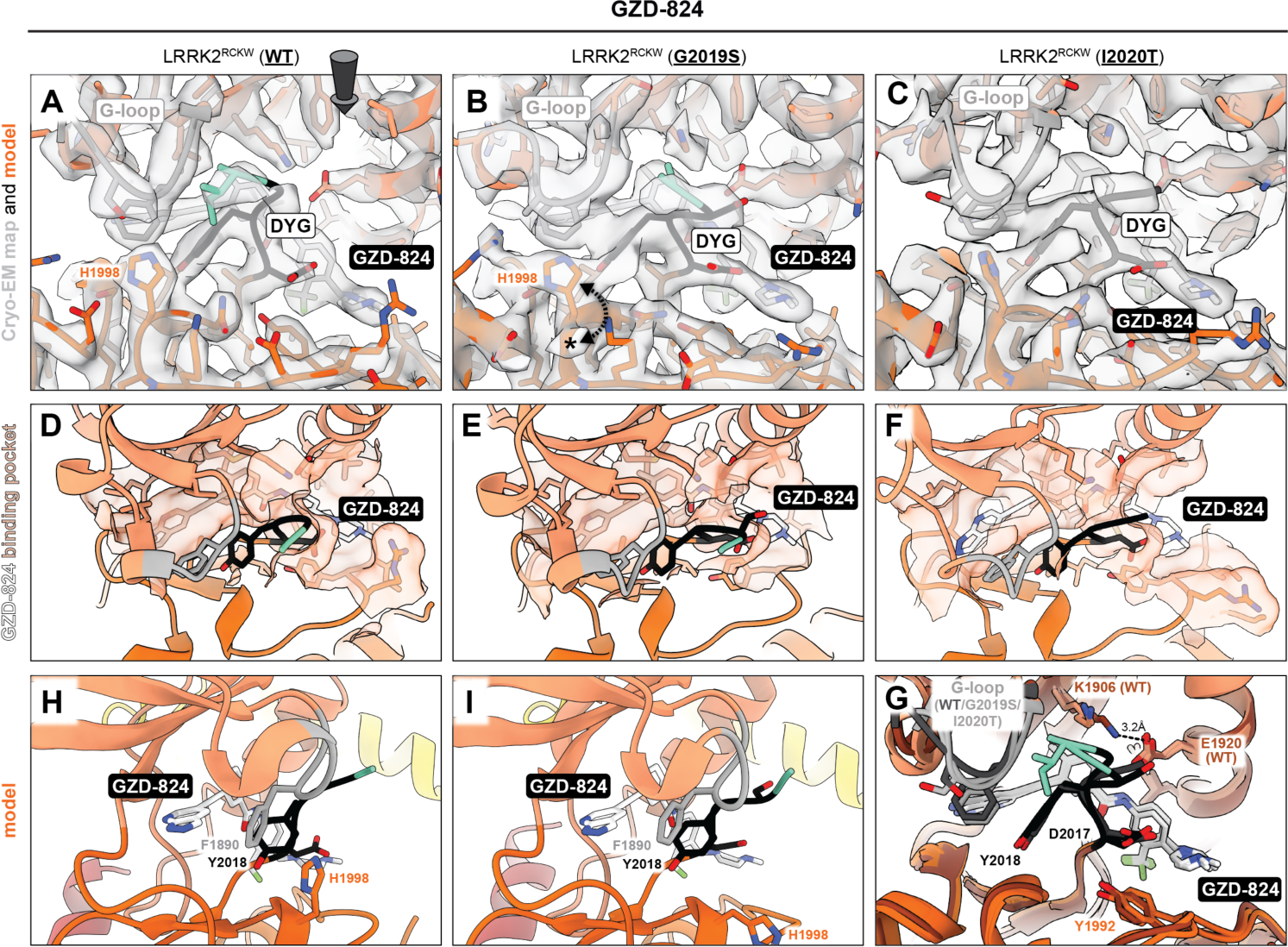
Binding of GZD-824 to LRRK2^RCKW^ wild-type (WT), G2019S, and I2020T. **(A,C)** Cryo-EM maps and models for the GZD-824 binding site and surrounding regions for LRRK2^RCKW^(WT):GZD-824 (A), LRRK2^R^^CKW^(G2019S):GZD-824 (B), and LRRK2^RCKW^(I2020T):GZD-824 The asterisks in (B) highlights a density present in G2019S but absent in WT. The dashed arrow in (B) indicate that two different rotamers of H1998 can be accommodated by the map of G2019S. The rotamer that points “up” (towards the N-lobe of the kinase) is shown in panel (B). Note that only density accounting for the “up” rotamer is seen in WT (A). The G-loop and DYG motif are indicated in the panels. The arrow in panel (A) indicates the viewing direction for panels (D-F). **(D-F)** Inhibitor binding pocket. The semitransparent orange surface indicates the solvent accessible surface for those residues in contact with GZD-824. **(G)** Superposition of the models for GZD-824 bound to LRRK2^RCKW^ WT, G2019S, and I2020T. WT is shown in darker shades. **(H,I)** Pi-pi interactions in the GZD-824 binding pocket. The GZD-824 binding pocket is viewed from the direction of the kinase’s hinge motif for WT (H) and G2019S (I) The G-loop’s F1890 faces the pyrazolopyridine ring of the inhibitor and, in turn, interacts with the Y2018 of the DYG motif. H1998 is parallel to F1890 in the WT structure, where it adopts the “up” rotamer (H). G2019S can accommodate two rotamers for H1998 (B), and the model in (I) shows H1998 in its “down” rotamer, which places it away from the inhibitor’s binding site.

As with the MLi-2-bound structures, we noticed differences in the rotamers adopted by H1998. In this case, however, the cryo-EM map for LRRK2^RCKW^(G2019S):GZD-824 could accommodate two rotamers for H1998 (**Figure 4B**), while WT (**Figure 4A**) showed density for a single rotamer. Although LRRK2^RCKW^(I2020T):GZD-824 appeared to also show a single rotamer (**Figure 4C**), this cryo-EM map was of lower resolution and more anisotropic, making it difficult to establish this side chain conformation unambiguously. A potential role for H1998 in the GZD-824-bound structures is in the stabilization of F1890 in the G-loop, which packed against the pyrazolopyridine ring of the inhibitor (**Figure 4H**). The presence of an alternative rotamer in G2019S (**Figure 4I**) could result in weaker binding of GZD-824 to this mutant. This prediction is consistent with a study showing that GZD-824 inhibits LRRK2(WT) in an *in vitro* phosphorylation assay with an IC_50_ ∼4-fold lower than that for LRRK2(G2019S) (*25*).

### Structures of full-length LRRK2 bound to MLi-2 and GZD-824

On cryo-EM grids, LRRK2 monomers were shown to adopt an autoinhibited conformation where the N-terminal repeats (ARM-ANK-LRR) sterically block access to the kinase’s active site (*13*). The repeats, which connect to the ROC domain, are held in place by an interaction between the “latch” helix, found within an insert in the LRR, and the WD40 domain (*13*). This anchoring of the repeats restricts conformational changes within LRRK2. To determine whether the presence of the repeats would prevent some of the conformational changes we observed in the structures we obtained with LRRK2^RCKW^, we solved structures of full-length LRRK2(I2020T) bound to either MLi-2 or GZD-824 (**Figure 5, Figures S9 and S10, Table S1**). As it was the case for all LRRK2^RCKW^ structures reported here, we solved the full-length ones in the presence of the E11 DARPin and GDP.

**Figure 5.**
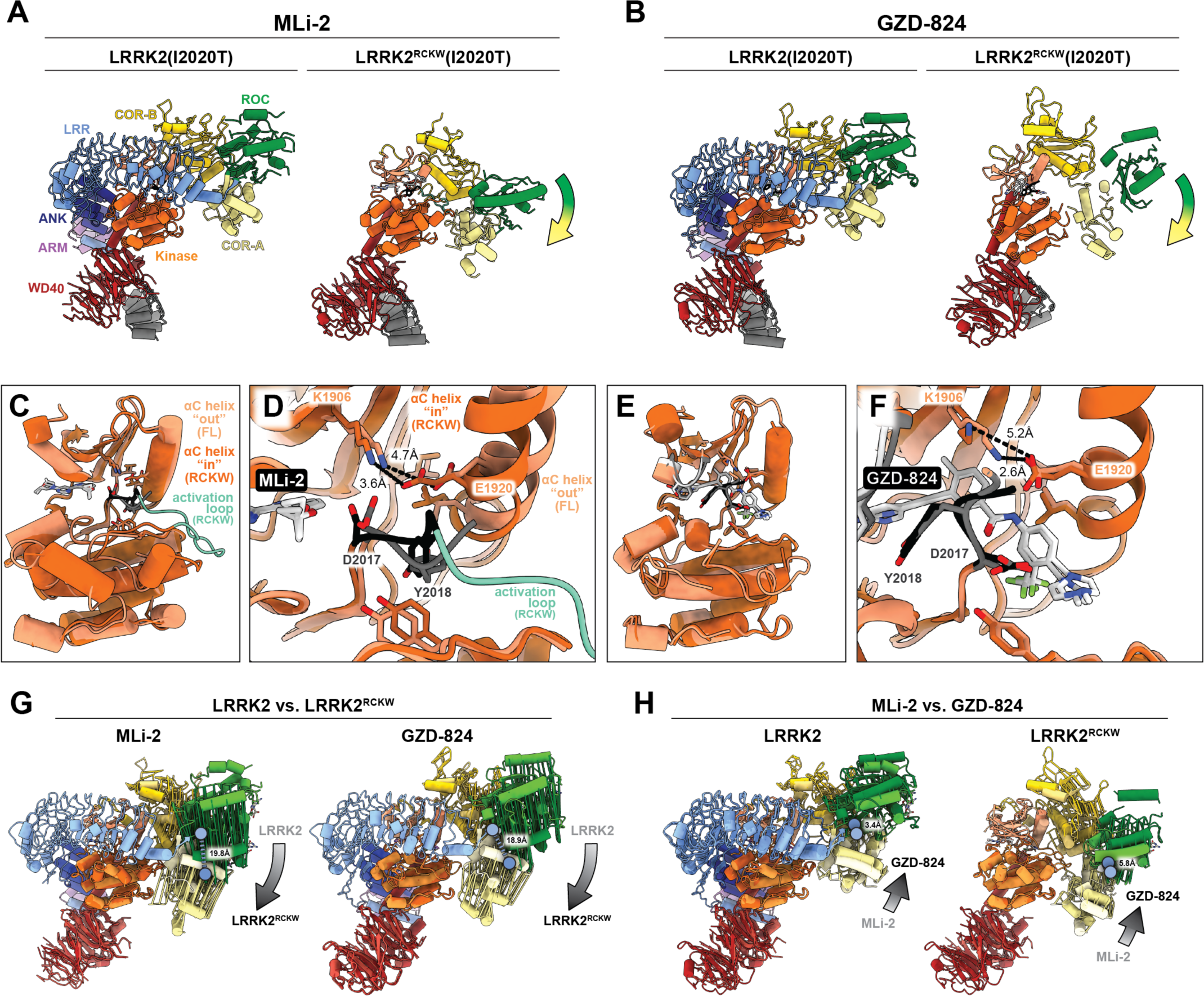
Structures of MLi-2 and GZD-824 bound to full-length LRRK2(I2020T) **(A)** Structures of LRRK2(I2020T) (left) and LRRK2^RCKW^(I2020T) (right) bound to MLi-2. (The structure of LRRK2^RCKW^(I2020T):MLi-2 is the same one shown in **Figure 3**.) The colored arrow indicates the movement of the ROC-COR domains in LRRK2^RCKW^ relative to full-length LRRK2. **(B)** Same as in (A), for GZD-824. (The structure of LRRK2^RCKW^(I2020T):GZD-824 is the same one shown in **Figure 4**.) **(C)** Overlay of the kinase domains from LRRK2(I2020T) and LRRK2^RCKW^(I2020T) bound to MLi-2, highlighting the “out” conformation of the αC helix. In panels (C) to (F), lighter shades represent LRRK2 and darker shades represent LRRK2^RCKW^. **(D)** Close-up of the kinase active sites in (C), with major features labeled. **(E)** Overlay of the kinase domains from LRRK2(I2020T) and LRRK2^RCKW^(I2020T) bound to GZD-824, highlighting their similarity. **(F)** Close-up of the kinase active sites in (C), with major features labeled. **(G)** Conformational differences between LRRK2 and LRRK2^RCKW^ bound to either MLi-2 (left) or GZD-824 (right). Structures were aligned using the C-lobes of their kinase domains. Colored vectors connect the same α carbons between the two structures. The gradient arrows highlight the direction of the overall relative movement of the ROC-COR moiety. The blue circles mark M1335 at the beginning of the ROC domain; the distance separating the α carbons of M1335 in the two structures being compared is indicated. **(H)** Differences in the conformations adopted by LRRK2 (left) or LRRK2^RCKW^ (right) in the presence of either MLi-2 or GZD-824. All labels and indicators are the same as those in (G).

As expected from the anchoring effect of the repeats, both LRRK2(I2020T):MLi-2 and LRRK2(I2020T):GZD-824 had their ROC-COR domains in a significantly more open (i.e. further away from the kinase) conformation than that seen in the equivalent structures with LRRK2^RCKW^ (**Figure 5A,B**). Next, we compared the kinase domains to see whether this difference in global conformation reflected differences in the states of the kinases themselves between LRRK2 and LRRK2^RCKW^ (**Figure 5C-F**). This was the case for MLi-2: while our structure of LRRK2^RCKW^(I2020T):MLi2 showed an active-like, DYG “in”, αC “in” conformation with an ordered activation loop, the kinase in LRRK2(I2020T):MLi-2 had its αC in an “out” conformation and a disordered activation loop. This difference, in contrast, was absent from the GZD-824 structures; the kinases in LRRK2(I2020T):GZD-824 and LRRK2^RCKW^(I2020T):GZD-824 were almost identical, the only difference being the formation of the salt bridge between K1906 and E1920 in LRRK2(I2020T):GZD-824 (**Figure 5E,F**).

To better understand this difference between MLi-2 and GZD-824, we compared the four structures carrying the I2020T mutation—LRRK2(I2020T):MLi-2, LRRK2^RCKW^(I2020T):MLi2, LRRK2(I2020T):GZD-824, and LRRK2^RCKW^(I2020T):GZD-824—by aligning them using the C-lobe of their kinases (**Figure 5G,H**). We used vectors connecting equivalent α carbons to illustrate the conformational changes, and we quantified differences between structures by measuring the distance between the α carbons of M1335, a residue located at the beginning of the ROC domain (**Figure 5G,H**). This analysis suggested that the repeats may be preventing the kinase in LRRK2(I2020T):MLi-2 from reaching its fully closed / active-like conformation.

While both MLi-2 and GZD-824 led to relatively similar conformational differences between LRRK2 and LRRK2^RCKW^, MLi-2 did seem to result in a slightly more closed conformation in LRRK2^RCKW^ (0.9Å between the α carbons of M1335) (**Figure 5G**), although we cannot determine whether this difference is significant at this point. The differences between the two types of inhibitors increased when we compared LRRK2(I2020T):MLi-2 to LRRK2(I2020T):GZD-824, and LRRK2^RCKW^(I2020T):MLi-2 to LRRK2^RCKW^(I2020T):GZD-824 (**Figure 5H**). As expected, the type-I inhibitor MLi-2 led to a more closed kinase conformation in LRRK2 than the type-II GZD-824 (**Figure 5H**, left); the ROC domain in the MLi-2 structure is shifted towards the kinase by 3.4Å as measured by the distance between the M1335 residues. This difference increased to 5.8Å in LRRK2^RCKW^, where the N-terminal repeats are absent(**Figure 5H**, right). Taken together, these observations suggested that the inability of MLi-2 to induce a fully closed / active-like state in the kinase of LRRK2(I2020T):MLi-2 was a result of the conformational constraining brought about by the anchored N-terminal repeats. Although we do not yet understand, mechanistically, what physiological processes lead to the release of the N-terminal repeats of LRRK2, and thus to full activation of the kinase, our work shows that MLi-2 was able to interact with the kinase both in the active-like state (LRRK2^RCKW^) and in an intermediate activation state (LRRK2) (**Figure 5C,D**).

## Discussion

Here, we have presented cryo-EM structures of LRRK2 and LRRK2^RCKW^ bound to the LRRK2-specific type-I inhibitor MLi-2, and the broad-spectrum type-II inhibitor GZD-824. These structures revealed the active-like state of LRRK2’s kinase (in the case of MLi-2), and the structural differences involved in engaging the two main types of inhibitors.

By comparing LRRK2^RCKW^(WT) with variants carrying the PD-linked mutations G2019S and I2020T, both of which are found within the kinase catalytic domain, we have begun to understand how some of these mutants might affect inhibitor binding. Similarly, comparing structures of inhibitor-bound LRRK2 and LRRK2^RCKW^ provided insights into how the N-terminal repeats of LRRK2 impact the interactions with inhibitors by constraining its conformational flexibility. These points are discussed further below.

### The role of the G2019S mutation in LRRK2 inhibition

A comparison of our structures of LRRK2^RCKW^(WT, G2019S, and I2020T) bound to MLi-2 showed a difference in the conformations that H1998, located in the C-lobe, could adopt (**Figure 3**). The cryo-EM maps of WT and I2020T showed density that could be accounted for by two different rotamers of H1998, while G2019S could only accommodate one. Importantly, one of the rotamers in WT and I2020T, the one absent in G2019S, contributed to a larger MLi-2-binding interface that closed around the inhibitor. The absence of this additional interaction in G2019S suggested that MLi-2 might bound to G2019S with lower affinity, a prediction that was consistent with our DSF measurements of LRRK2^KW^ upon addition of MLi-2 (**Figure 1**).

A paper from the West group reported that the G2019S mutation made LRRK2 resistant to inhibition by MLi-2 *in vivo*. Although this would appear to agree with our structural and functional data, this study relied on measuring the loss of LRRK2-S935 phosphorylation as a proxy for inhibition (*33*). While this is a frequently used biomarker for LRRK2 inhibition, it is a correlative one that is not directly dependent on LRRK2’s kinase activity. A direct measurement of LRRK2’s kinase activity, by quantifying the levels of Rab10 phosphorylation, was not possible in that study (*33*). On the other hand, data published by the Alessi group using purified LRRK2 and an *in vitro* assay with a peptide kinase substrate came to the opposite conclusion: that MLi-2 inhibited LRRK2(G2019S) ∼2-fold better than LRRK2(WT) (*25*). As we discuss below, we hypothesize that these discrepancies may reflect the type of construct used (full-length LRRK2 in the Alessi study, LRRK2^KW^ in ours), and the potential roles played by the N-terminal repeats of LRRK2 in conformational changes and inhibitor binding.

We also saw differences in the rotamers that H1998 could adopt in our structures of LRRK2^RCKW^ bound to GZD-824 (**Figure 4**). In this case, the cryo-EM map of LRRK2^RCKW^(G2019S):GZD-824 showed density indicative of two rotamers for H1998, only one of which would contribute to inhibitor binding, while we only saw the latter in the cryo-EM map of the WT complex. The prediction, based on this structural difference, that LRRK2(WT) might bind more tightly than G2019S to GZD-824 was consistent with *in vitro* measurements of the IC_50_ values for GZD-824 for these two variants (*25*). The G-loop adopted slightly different conformations in our LRRK2^RCKW^(WT):GZD-824 and LRRK2^RCKW^(G2019S):GZD-824 structures, and this could in principle have an effect on H1998, which is close to the G-loop in its “up” rotamer (**Figure 4A,H**). However, our model of LRRK2^RCKW^(I2020T):GZD-824 was very similar to the one of G2019S (**Figure 4G**), yet the corresponding cryo-EM map appeared to show density for a single H1998 rotamer, equivalent to that in WT. The caveat, which we noted earlier, is that the cryo-EM map for I2020T was of lower resolution and more anisotropic than those for WT and G2019S, so it was not possible at this point to rule out the possibility that the conformation of the G-loop was responsible for the difference between WT and G2019S mutant.

### The role of the N-terminal repeats in LRRK2 inhibition

The N-terminal repeats of LRRK2 appear to play a double role in autoinhibition. On the one hand, the LRR domain physically blocks access to the kinase’s active site (*13*). On the other, the LRR restricts the conformational flexibility of the C-terminal half of LRRK2 (LRRK2^RCKW^), to which it is connected at the ROC domain, by anchoring itself to the WD40 domain via its “latch” helix (*13*). Our analysis of the conformational differences between LRRK2 and LRRK2^RCKW^ bound to MLi-2 and GZD-824 (**Figure 5**) suggested that the constrains imposed by the N-terminal repeats had a bigger effect on type-I inhibitors. While our structures of LRRK2^RCKW^:MLi-2 (**Figures 2 and 3**) showed the kinase in a fully closed / active-like state, as would be expected from a type-I inhibitor, the kinase in LRRK2(I2020T):MLi-2 was only partially closed, with its αC “out” and a disordered activation loop. We attribute this intermediate state to the tug-of-war between the MLi-2-induced closing of the kinase, and the constraining effect of the N-terminal repeats. We did not see this difference between LRRK2(I2020T):GZD-824 and LRRK2^RCKW^(I2020T):GZD-824; the kinases in the two structures, in an inactive / open conformation, were almost identical (**Figure 4**).

Taken together, these comparisons suggest that the N-terminal repeats of LRRK2 prevented LRRK2 from reaching the fully closed / active-like state. At the same time, our structures showed that MLi-2 can bind to LRRK2 regardless of whether the kinase is fully closed (LRRK2^RCKW^) or in an intermediate activation state (LRRK2). Our results raise the possibility that IC_50_’s measured using full-length LRRK2 (*in vitro*, in cells or *in vivo*) may be a combination of IC_50_’s for at least two different populations: one population corresponds to the autoinhibited LRRK2, where the kinase cannot reach its fully active-like state, represented by our LRRK2(I2020T):MLi-2 structure; the other population corresponds to activated LRRK2, where the repeats have undocked and the kinase can reach its active-like state, represented by our LRRK2^RCKW^(G2019S):MLi-2 structure.

The structures presented here provide a blueprint for medicinal chemistry efforts to design new LRRK2-specific inhibitors, in particular type-II inhibitors for which we have no example of LRRK2 selective compounds. As mentioned in the introduction, selective type-II inhibitors are needed not only to expand the potential therapeutic arsenal to treat PD, but also to create a toolkit that allows inhibition LRRK2 in conformation-specific states to better understand the function of this protein in cells and *in vivo*.

## Acknowledgements

We would like to thank Samara Reck-Peterson for feedback on the manuscript. We also thank the UC San Diego Cryo-EM Facility, and the Physics Computing Facility at UC San Diego for IT support.

## Author contributions

Conceptualization: SM, SK, AEL

Preparation of reagents: MSM, AVS, VD, DC, JAL, SM

Data collection & analysis: MSM, AVS, VD, DC, JAL

Supervision: SM, SK, AEL

Writing & editing: MSM, AVS, SM, SK, AEL

## Data and materials availability

Raw data for the assays shown in **Figures 1, 3H, and S1** have been uploaded to Zenodo (DOI: https://zenodo.org/record/8323739). Cryo-EM maps and associated models have been deposited in the EM Data Bank and Protein Data Bank, respectively. Accession codes are as follows: LRRK2^RCKW^(WT):MLi-2:E11 (tetramer) (EMDB: 41709; PDB: 8TXZ); LRRK2^RCKW^(G2019S):MLi-2:E11 (tetramer) (EMDB: 41754; PDB: 8TZC); LRRK2^RCKW^(I2020T):MLi-2:E11 (tetramer) (EMDB: 41758; PDB: 8TZG); LRRK2^RCKW^(WT):GZD-824:E11 (trimer) (EMDB: 41756; PDB: 8TZE); LRRK2^RCKW^(G2019S):GZD-824:E11 (monomer) (EMDB: 41728, 41797, 41798, and 41799; PDB: 8TYQ); LRRK2^RCKW^(I2020T):GZD-824:E11 (monomer) (EMDB: 41753, 41794, 41795, and 41802; PDB: 8TZB); LRRK2(I2020T):MLi-2:E11 (monomer) (EMDB: 41759; PDB: 8TZH); and LRRK2(I2020T):MLi-2:E11 (monomer) (EMDB: 41757; PDB: 8TZF). All other data are available in the main text or the supplementary materials.

## Materials and Methods

### Cloning, plasmid construction, and mutagenesis

Briefly, the DNA coding for LRRK2-WT residues 1327 to 2527 (taken from Mammalian Gene Collection) was PCR-amplified using the forward primer TACTTCCAATCCATGAAAA AGGCTGTGCCTTATAACCGA and the reverse primer TATCCACCTTT ACTGTCAC-TCAACAGATGTTCGTCTCATTTTTTCA. The amplicon was then inserted into the expression vector pFB-6HZB by ligation-independent cloning. This plasmid was used as a template for G2019S and I2020T site-directed mutagenesis (Q5 Site-Directed Mutagenesis Kit, NEB, Catalog # E0445S). These plasmids were used for the generation of recombinant baculoviruses following Bac-to-Bac expression system protocols (Invitrogen, Catalog # 10359016).

### Inhibitors

Stocks of the kinase inhibitors MLi-2 (10 mM; Tocris) and GZD-824 (10 mM; Cayman Chemical), were stored in DMSO at -20 C.

### Dynamic scanning fluorimetry (DSF) assay

The assay was performed according to a previously established protocol (*34*). Briefly, a solution of 4 µM LRRK2^KW^ protein in assay buffer (20 mM HEPES pH 7.4, 150 mM NaCl, 0.5 mM TCEP, 20 µM GDP, 2.5 mM MgCl_2_, 0.5% glycerol) was mixed 1:1000 with SYPRO Orange (Sigma). The inhibitors to be tested were added to a final concentration of 10 µM. 20 µL of each sample were placed in a 96-well plate and heated gradually from 25°C to 95°C. Fluorescence was monitored using an Mx3005P real-time PCR instrument (Stratagene) with excitation and emission filters set to 465 and 590 nm, respectively. Data was analysed with the MxPro (https://www.genomics.agilent.com/article.jsp?pageId=2100; RRID:SCR_016375) software. Protocols are also available in protocols.io (DOI: dx.doi.org/10.17504/protocols.io.kxygx3y6kg8j/vi).

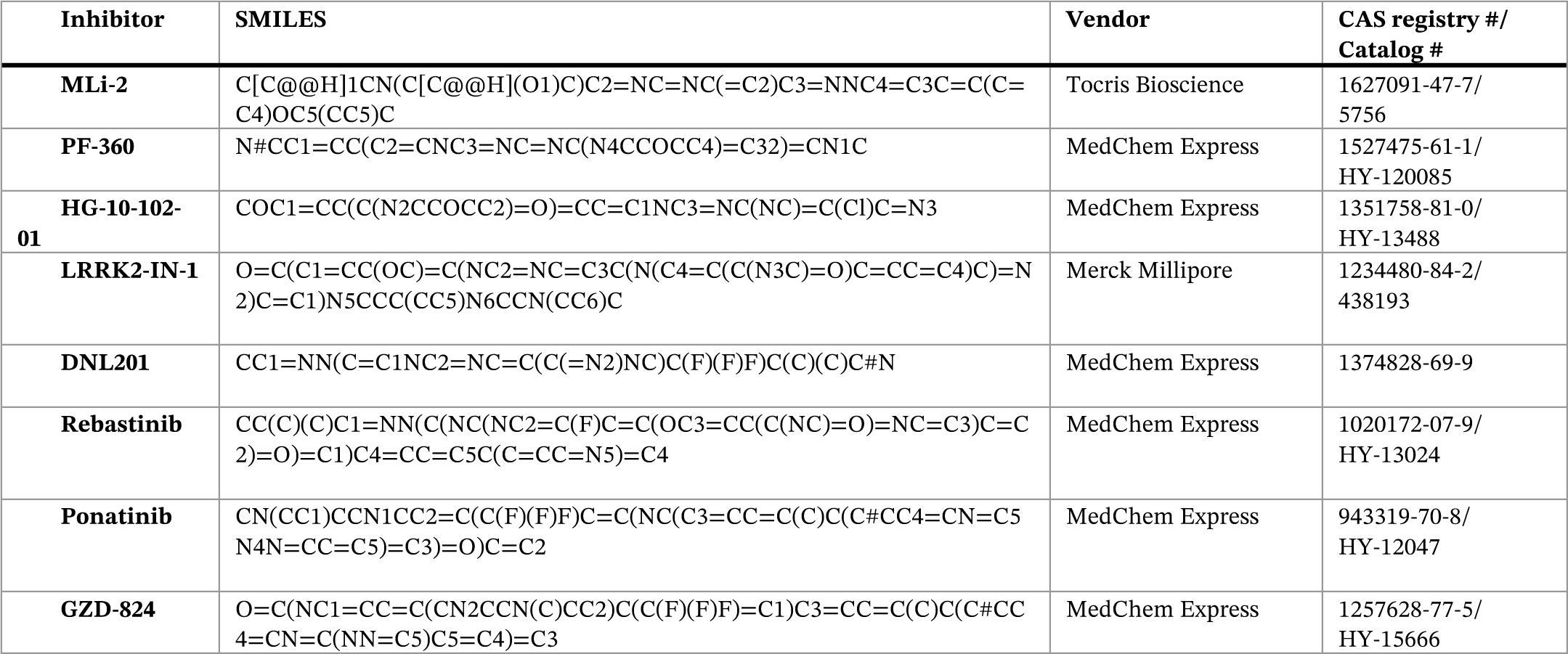

### Mass spectroscopy (MS)-based activity assay

To determine the IC_50_ values for a set of LRRK2 inhibitors, the phosphorylation of the LRRK2 substrate Rab8A was assessed applying an MS-based activity assay. 50 nM LRRK2^RCKW^ protein was mixed with 5 µM substrate and varying concentrations of the inhibitors in assay buffer (20 mM HEPES pH 7.4, 150 mM NaCl, 0.5 mM TCEP, 20 µM GDP, 2.5 mM MgCl_2_, 0.5% glycerol). The reaction was started by adding 1 mM ATP and incubated at room temperature. The reaction was stopped by adding MS buffer (0.1 % formic acid in water) and subjected to an Agilent 6230 Electrospray Ionization Time-of-Flight mass spectrometer coupled with the liquid chromatography unit 1260 Infinity for analysis. 5 µL of the reaction mix was injected onto a C3 column and eluted at 0.4 mL/min flow rate using a solvent gradient of water to acetonitrile with 0.1% formic acid. Data was acquired using the MassHunter LC/MS Data Acquisition software and analyzed using the BioConfirm vB.08.00 tool (both Agilent Technology). The peak intensities of the unphosphorylated and phosphorylated Rab8A species were quantified and the relative kinase activity calculated. To determine the IC_50_ values, a non-linear regression with variable slope was fitted to the datapoints with GraphPad Prism (http://www.graphpad.com; RRID:SCR_002798). Protocols are also available in protocols.io (DOI: dx.doi.org/10.17504/protocols.io.6qpvr385ovmk/v2).

### Protein purification: LRRK2^RCKW^ and LRRK2

For both LRRK2^RCKW^ and LRRK2 purifications, the pelleted Sf9 cells were washed with PBS, resuspended in lysis buffer (50 mM HEPES pH 7.4, 500 mM NaCl, 20 mM imidazole, 0.5 mM TCEP, 5% glycerol, 5 mM MgCl_2_, 20 µM GDP) and lysed by homogenization. The supernatant was cleared by centrifugation and loaded onto a Ni-NTA (Qiagen) column. After rinsing with lysis buffer, the His_6_-Z-tagged protein was eluted in lysis buffer containing 300 mM imidazole. The eluate was then diluted to 250 mM NaCl mM with dilution buffer (50 mM HEPES pH 7.4, 0.5mM TCEP, 5% glycerol, 5mM MgCl_2_, 20µM GDP) and loaded onto an SP sepharose column. His_6_-Z-TEV-LRRK2^RCKW^/His_6_-Z-TEV-LRRK2 were eluted with a gradient of 250 mM to 2.5 M NaCl in dilution buffer and then treated with TEV protease overnight to cleave the His_6_-Z. Contaminating proteins, the cleaved tag, uncleaved protein and TEV protease were removed by another combined SP sepharose Ni-NTA step. Finally, LRRK2^RCKW^ was concentrated and subjected to gel filtration in 20 mM HEPES pH 7.4, 700 mM NaCl, 0.5 mM TCEP, 5% glycerol, 2.5 mM MgCl_2_, 20 µM GDP, while LRRK2 was concentrated and subjected to gel filtration in 20 mM HEPES pH 7.4, 200 mM NaCl, 0.5 mM TCEP, 5% glycerol, 2.5 mM MgCl_2_, 20 µM GDP using an AKTA Xpress system with an S200 gel filtration column. Final yields, as calculated from UV absorbance, are typically 1.2-3.3 mg of LRRK2^RCKW^ and 0.9-2.2 mg of LRRK2 per liter of insect cell medium. Protocols are also available in protocols.io for the expression of LRRK2 (DOI: dx.doi.org/10.17504/protocols.io.rm7vzyyrrlx1/v1), and the purification of LRRK2^RCKW^ (DOI: dx.doi.org/10.17504/protocols.io.81wgb6693lpk/v1), and full-length LRRK2 (DOI: dx.doi.org/10.17504/protocols.io.rm7vzx3b5gx1/v1).

### Cryo-EM sample preparation for LRRK2^RCKW^:MLi2/GZD-824:E11 DARPin complexes

Purified LRRK2^RCKW^ (WT, G2019S or I2020T) was exchanged into 20 mM HEPES pH 7.4, 150 mM NaCl, 0.5mM TCEP, 5% glycerol, 2.5mM MgCl_2_ and 20µM GDP. LRRK2^RCKW^ was incubated with E11-DARPin in a 1:1.25 molar ratio and either MLi-2 (20 µM) or GZD-824 (40 µM) for 10 minutes at RT and 15 minutes at 4°C. Complexes were diluted to a final concentration range of 4 to 6 µM in the same buffer before plunge freezing. 3 or 3.5 μl of LRRK2^RCKW^:inhibitor:E11 DARPin complex were applied to a glow-discharged UltraAuFoil Holey Gold 200 mesh R2/2 grid (Quantifoil, Catalog # Q250AR2A) and incubated in a FEI Vitrobot IV chamber at 4°C and 95% humidity for 20 sec. The excess liquid was blotted for 4 sec using filter paper 595 (Ted Pella, Prod # 47000-100) at blot force 3, and vitrified by plunging into liquid ethane cooled down to liquid nitrogen temperature. Protocol is also available in protocols.io: DOI: dx.doi.org/10.17504/protocols.io.q26g7p17qgwz/v1.

### Cryo-EM sample preparation for full length LRRK2(I2020T):MLi-2/GZD-824-E11 DARPin complex

Purified full-length LRRK2(I2020T) was incubated with E11 DARPin at a molar ratio of 1:1.25 and either MLi-2 (20 μM) or GZD-824 (40 μM) for 10 minutes at RT and 5-10 minutes at 4°C. Complexes were diluted to a 5 µM final concentration in the same buffer (20 mM HEPES pH 7.4, 150 mM NaCl, 0.5 mM TCEP, 5% glycerol, 2.5 mM MgCl_2_) with the addition of a final 20 µM GDP for LRRK2(I2020T):MLi-2:E11, or 100 µM GMP-PNP for LRRK2(I2020T):GZD-824:E11. 3 to 3.5 μl of complex was applied to a glow-discharged UltrAuFoil Holey Gold 200 mesh R2/2 grid (Quantifoil, Catalog # Q250AR2A) and incubated in a FEI Vitrobot IV chamber at 4°C and 95% humidity for 20 sec. The excess liquid was blotted for 4 sec using filter paper 595 (Ted Pella, Prod # 47000-100) at blot force 4, vitrified by plunging into liquid ethane cooled down to liquid nitrogen temperature.

### Cryo-EM data collection

Cryo-EM data for LRRK2^RCKW^(G2019S and I2020T):MLi-2:E11 DARPin, LRRK2^RCKW^(WT, G2019S and I2020T):GZD-824:E11 DARPin and LRRK2(I2020T):MLi-2:E11 DARPin were collected on a Titan Krios G3 (Thermo Fisher Scientific) operated at 300 keV, equipped with a Falcon 4 direct electron detector (Thermo Fisher Scientific) and a Gatan BioContinuum energy filter. Images were collected at a nominal magnification of 130,000x in EF-TEM mode (0.935 Å calibrated pixel size) using a 20-eV slit width in the energy filter and a cumulative electron exposure of ∼55 electrons/Å^2^. Data for LRRK2(I2020T)-E11:GZD-824 was collected on a Talos Arctica (FEI) operated at 200 keV, equipped with a Falcon 4i (Thermo Fisher Scientific) with a cumulative electron exposure of ∼55 electrons/Å^2^. Data were collected automatically using EPU software (Thermo Fisher Scientific). Data for LRRK2^RCKW^:MLi-2:E11 DARPIn were collected on a Titan Krios G3 (Thermo Fisher Scientific) operated at 300 keV, equipped with a K3 Summit direct electron detector (Gatan) and a Gatan BioContinuum energy filter. Images were collected at a nominal magnification of 105000x in EF-TEM mode (0.822 Å calibrated pixel size) using a 20-eV slit width in the energy filter and a cumulative electron exposure of approximately 57 electrons/Å^2^.

### Data processing for LRRK2_RCKW_:MLi2:E11 DARPin complexes

11,181, 8,402 and 10,488 movies were collected for LRRK2^RCKW^-WT:MLi-2:E11 DARPin, LRRK2^RCKW^(G2019S):MLi-2:E11 DARPin and LRRK2^RCKW^(I2020T):MLi-2:E11 DARPIn complexes, respectively. They were aligned using MotionCor2 (https://emcore.ucsf.edu/cryoem-software; RRID:SCR_016499) dose-weighted alignment option and CTF parameters were estimated on dose-weighted images using CTFFIND4 (http://grigorief-flab.janelia.org/ctffind4; RRID:SCR_016732) (*35*). Micrographs with a CTF fit worse than 3.5-4Å (as determined by CTFFIND4 (http://grigorief-flab.janelia.org/ctffind4; RRID:SCR_016732), and varying among datasets) were discarded from further processing. Particles were picked using a Topaz model previously trained for each dataset. Several rounds of reference-free 2D classification yielded a stack of 574,664 (WT), 578,873 (G2019S) and 694,336 particles (I2020T), each containing both monomers and tetramers. All particles were extracted with a 400-pixel box using CryoSPARC (https://cry-osparc.com; RRID:SCR_016732) (*36*) and were separated based on oligomerization state (**Supplementary Figure S2, S5 and S6**). Subsequently, ab-initio jobs were run to further remove bad particles. The best ab-initio volume for the tetramer was used as an input for NU-Refinement (D2 symmetry). Then, particles were expanded based on the volume symmetry and used in subsequent jobs. After 3D Variability and C1-symmetry local refinement, we obtained maps at 3.05, 2.74 and 2.8Å, respectively. The best ab-initio map for the monomer, along with monomer particles (**Figure S2**, S5 and S6) were used as an input for a NU-Refinement. In all maps, FSC and local resolution estimations were performed using the routines implemented in CryoSPARC (https://cry-osparc.com; RRID:SCR_016732) (*36*). Protocol is also available in protocols.io: DOI: dx.doi.org/10.17504/protocols.io.n2bvj3nmnlk5/v1.

### Data processing for LRRK2(I2020T):MLi-2:E11 DARPin complex

10,407 movies were collected for LRRK2(I2020T):MLi-2:E11 DARPin and preprocessed as previously described. Micrographs with a CTF fit worse than 4Å (as determined by CTFFIND4, http://grigoriefflab.janelia.org/ctffind4; RRID:SCR_016732) were discarded from further processing. Particle picking was done following the same protocol for the LRRK2^RCKW^ datasets. Several rounds of 2D classification yielded 162,409 particles. After Ab-initio job and a NU-Refinement, a final map at a resolution of 3.9Å was obtained (**Supplementary Figure S9**). Protocol is also available in protocols.io: DOI: dx.doi.org/10.17504/protocols.io.dm6gp3j4dvzp/v1.

### Data processing for LRRK2^RCKW^:GZD-824:E11 DARPin complexes

5,565, 7,988 and 8,386 movies were collected for LRRK2^RCKW^WT:GZD-824:E11 DARPin, LRRK2^RCKW^(G2019S):GZD-824:E11 DARPin and LRRK2^RCKW^(I2020T):GZD-824:E11 DARPin complexes, respectively. They were aligned using MotionCor2 (https://emcore.ucsf.edu/cryoem-software; RRID:SCR_016499) dose-weighted alignment option and CTF parameters were estimated on dose-weighted images using CTFFIND4 (http://grigorief-flab.janelia.org/ctffind4; RRID:SCR_016732) (*35*). Micrographs with a CTF fit worse than 3.5-4.5 Å (as determined by CTFFIND4, http://grigorief-flab.janelia.org/ctffind4; RRID:SCR_016732, and varying among datasets) were discarded from further processing. Particles were picked using a Topaz model previously trained for each dataset. Several rounds of reference-free 2D classification yielded a stack of 249,808, 759,293 and 735,633 monomer particles respectively; additionally, LRRK2^RCKW^WT:GZD-824:E11 dataset also yielded 58,678 trimer particles. Particles were extracted with a 320-pixel box for monomers, and 400-pixel box for trimers using CryoSPARC (https://cryosparc.com; RRID:SCR_016732) (*36*) (**Supplementary Figure S3**, S7 and S8). For all LRRK2^RCKW^ monomer particles, ab-initio jobs followed by heterogeneous refinement were run to sort particles. The best refined class was subject to a NU-refinement (C1 symmetry), followed by a local refinement with a mask surrounding the kinase-WD40-DARPin, yielding maps at resolutions of 3.10Å, 2.99Å, and 3.06Å, respectively. To account for the heterogeneity of the ROC-COR domain a 3D Variability job was done to further split the particles, followed by a local refinement with a mask surrounding the ROC-COR domain; this yielded maps at resolutions of 3.80Å, 3.50Å, and 3.63Å, respectively. For LRRK2^RCKW^ (trimer particles) an ab-initio job was run followed by a NU-refinement (C3 symmetry). Particles were expanded based on the volume symmetry and used in a C1-symmetry local refinement, yielding a map at 2.90Å resolution. Protocol is also available in protocols.io: DOI: dx.doi.org/10.17504/proto-cols.io.81wgbxz81lpk/v1.

### Data processing for LRRK2(I2020T):GZD-824:E11 DARPin complexes

4,102 movies were collected for LRRK2(I2020T):GZD-824:E11 and pre-processed as previously described. Micrographs with a CTF fit worse than 5.3Å (as determined by CTFFIND4, http://grigoriefflab.janelia.org/ctffind4; RRID:SCR_016732) were discarded from further processing. Particle picking was done in the same manner as for the LRRK2^RCKW^ datasets, followed by several rounds of 2D classification, resulting in 1,029,496 particles. Particles were moved to Relion (http://www2.mrc-lmb.cam.ac.uk/relion; RRID:SCR_016274) using pyem/csparc2star (https://github.com/asarnow/pyem/blob/master/csparc2star.py) and extracted to 3.8Å/pix. Particles were subject to 3D classification (5 classes) with alignment, 224,119 particles yielded the best 3D class of monomeric LRRK2(I2020T). Consensus 3D refinement using the best 3D class was done, followed by two rounds of 3D classification without alignment resulting in the best 3D class containing 60,465 particles. Particles were moved back to CryoSPARC (https://cryosparc.com; RRID:SCR_016732) (*34*), followed by a NU-refinement, yielding a map at 3.40Å resolution (**Supplementary Figure S10**). Protocol is also available in protocols.io: DOI: dx.doi.org/10.17504/protocols.io.dm6gp3j4dvzp/v1.

### Model building and refinement of LRRK2^RCKW^:MLi-2/GZD-824:E11 DARPin and LRRK2(I2020T):MLi-2/GZD-824:E11 DARPin

LRRK2^RCKW^ and LRRK2 models were built using the highest-resolution maps obtained for each complex. These maps corresponded to tetramers for LRRK2^RCKW^(WT):MLi-2:E11, LRRK2^RCKW^(G2019S):MLi-2:E11, and LRRK2^RCKW^(I2020T):MLi-2:E11, a trimer for LRRK2^RCKW^(WT):GZD-824:E11 and a monomer for FL(I2020T):MLi-2/GZD-824:E11 DARPin (see **Figures S2, S3, S5**-**S10** for details on the cryo-EM workflow for each specific sample). To rule out structural differences in the kinase induced by oligomerization, we tested the fit of the models built using the higher-resolution oligomers into cryo-EM maps of the monomeric form of the same construct (**Figure S4**).

Available PDBs 6VP7, 7LHW and A1N for LRRK2^RCKW^, LRRK2 and MLi-2, respectively, were used as starting points. Protein models were split into domains, docked into the corresponding cryo-EM maps, and merged. To obtain a GZD-824 model, electronic Ligand Building and Optimization Workbench (elBOW) software available in Phenix (https://www.phenix-online.org; RRID:SCR_014224) (*37*) was run using Simplified Molecular Input Line Entry System (SMILES) notation of molecule as an input. Inhibitor models (MLi-2 or GZD-824) were fitted in the map density and incorporated in the model. A combination of manual inspection of amino acids in Coot (http://www2.mrc-lmb.cam.ac.uk/personal/pemsley/coot; RRID:SCR_014222) (*38, 39*) and refinement of model into their maps in Phenix (https://www.phenix-online.org; RRID:SCR_014224) (*37*) was used to generate the final models.

### Analysis of conformational changes in LRRK2^RCKW^ and LRRK2

The segments linking equivalent α carbons in LRRK2^RCKW^ and LRRK2 were generated by aligning models by their C-lobes (residues 1949-2139) using UCSF ChimeraX (https://www.cgl.ucsf.edu/chimerax; RRID:SCR_015872) (*40*). Amino acids not present in the two models being compared were removed and primary sequences were aligned using ClustalW (EMBL-EBI)(*41*). Aligned sequences were used as an input for the public script PDBarrows.ipynb on Jupyter notebook, generating output arrows, which were colored to match the domains being linked. (**Figure 5 G,H**).

### Structure depictions

All structure-related figures were prepared using ChimeraX (https://www.cgl.ucsf.edu/chimerax; RRID:SCR_015872) (*40*).

---

**Figure S1.**
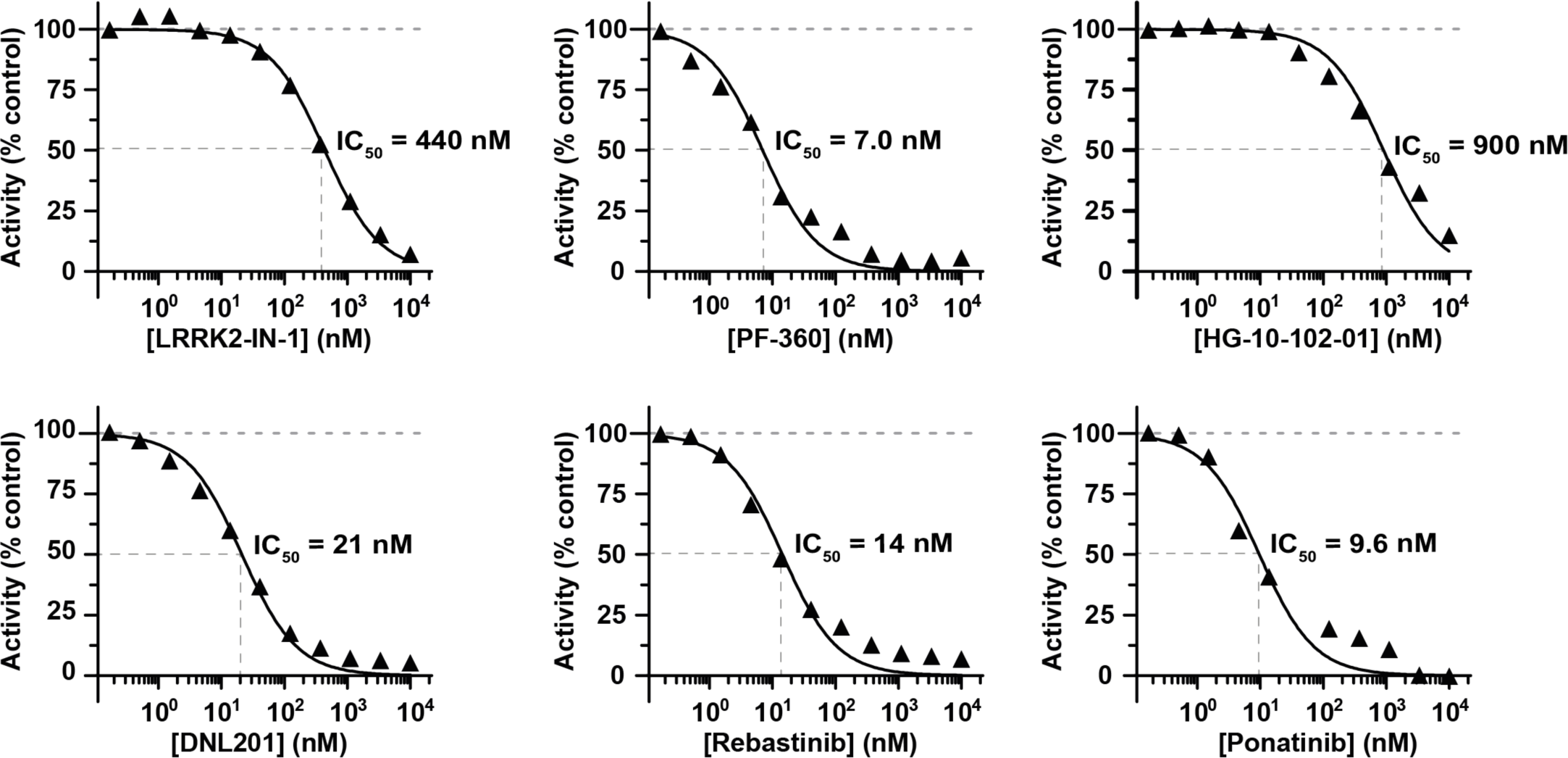
LRRK2 inhibition curves obtained from the MS-based activity assay. The physiological LRRK2 substrate Rab8A was subjected to phosphorylation by LRRK2^RCKW^. Varying concentrations of the indicated inhibitors were present in the reaction mixtures allowing for the determination of IC_50_ values.

**Figure S2.**
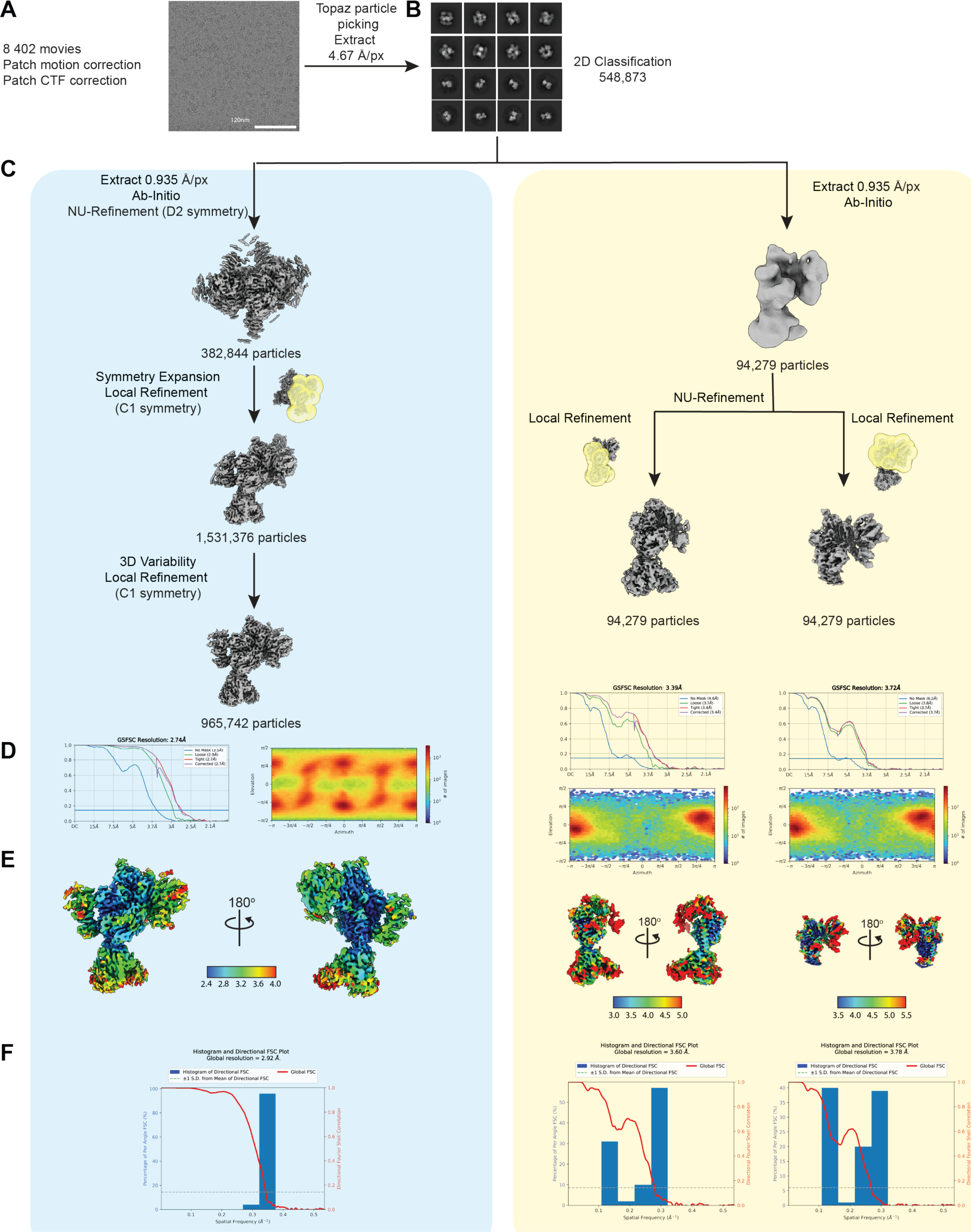
Cryo-EM workflow for LRRK2^RCKW^(G2019S):MLi-2. Representative micrograph **(A)**, 2D class averages **(B)**, data processing strategy **(C)**, FSC plots and Euler angle distributions **(D)**, local resolution maps **(E)**, and 3D FSC plot **(F)** for LRRK2^RCKW^(G2019S):MLi-2 tetramer (blue panel) and monomer (yellow panel).

**Figure S3.**
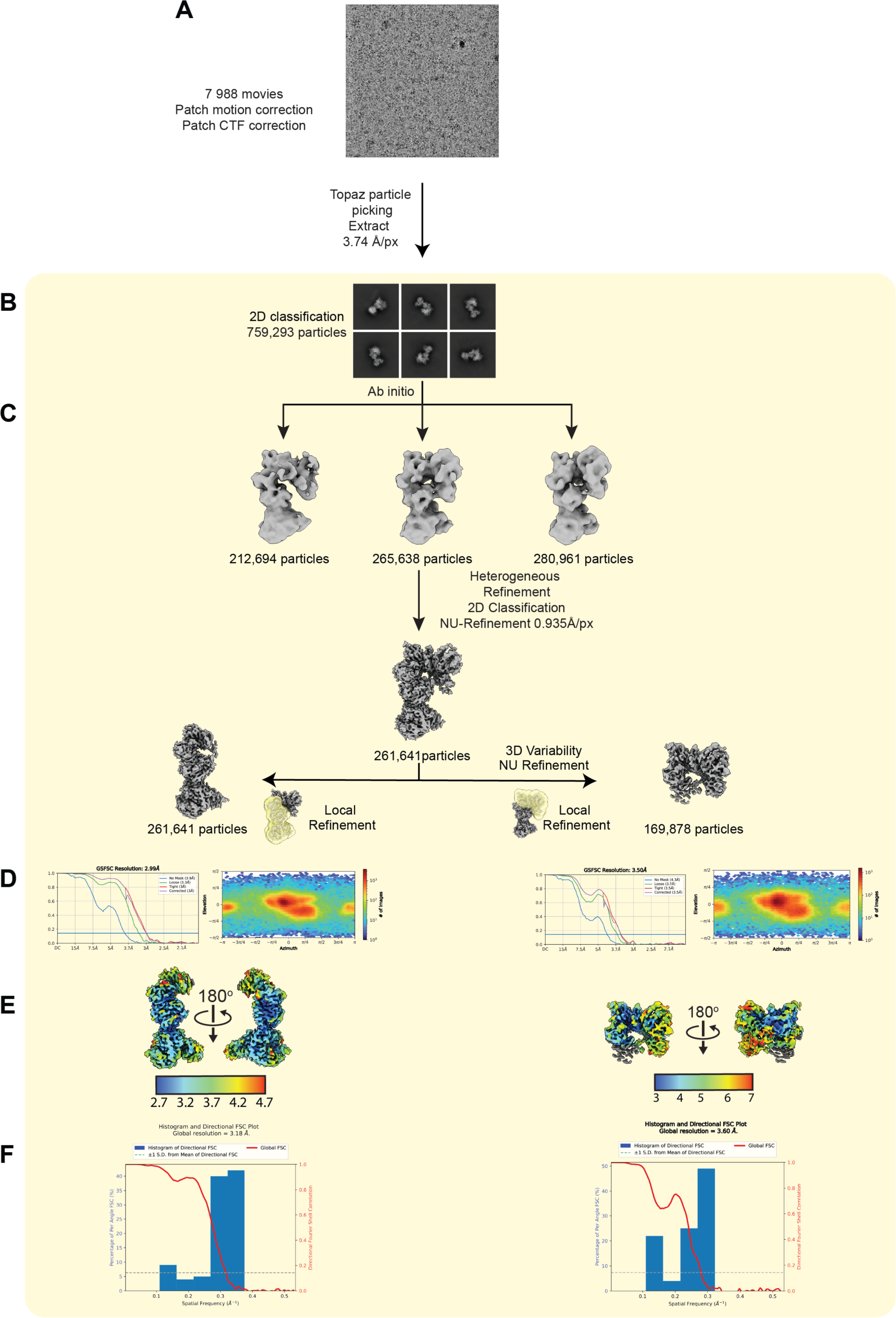
Cryo-EM workflow for LRRK2^RCKW^(G2019S):GZD-824. Representative micrograph **(A)**, 2D class averages **(B)**, data processing strategy **(C)**, FSC plots and Euler angle distributions **(D)**, local resolution maps **(E)**, and 3D FSC plot **(F)** for LRRK2^RCKW^(G2019S):GZD-824 monomer.

**Figure S4.**
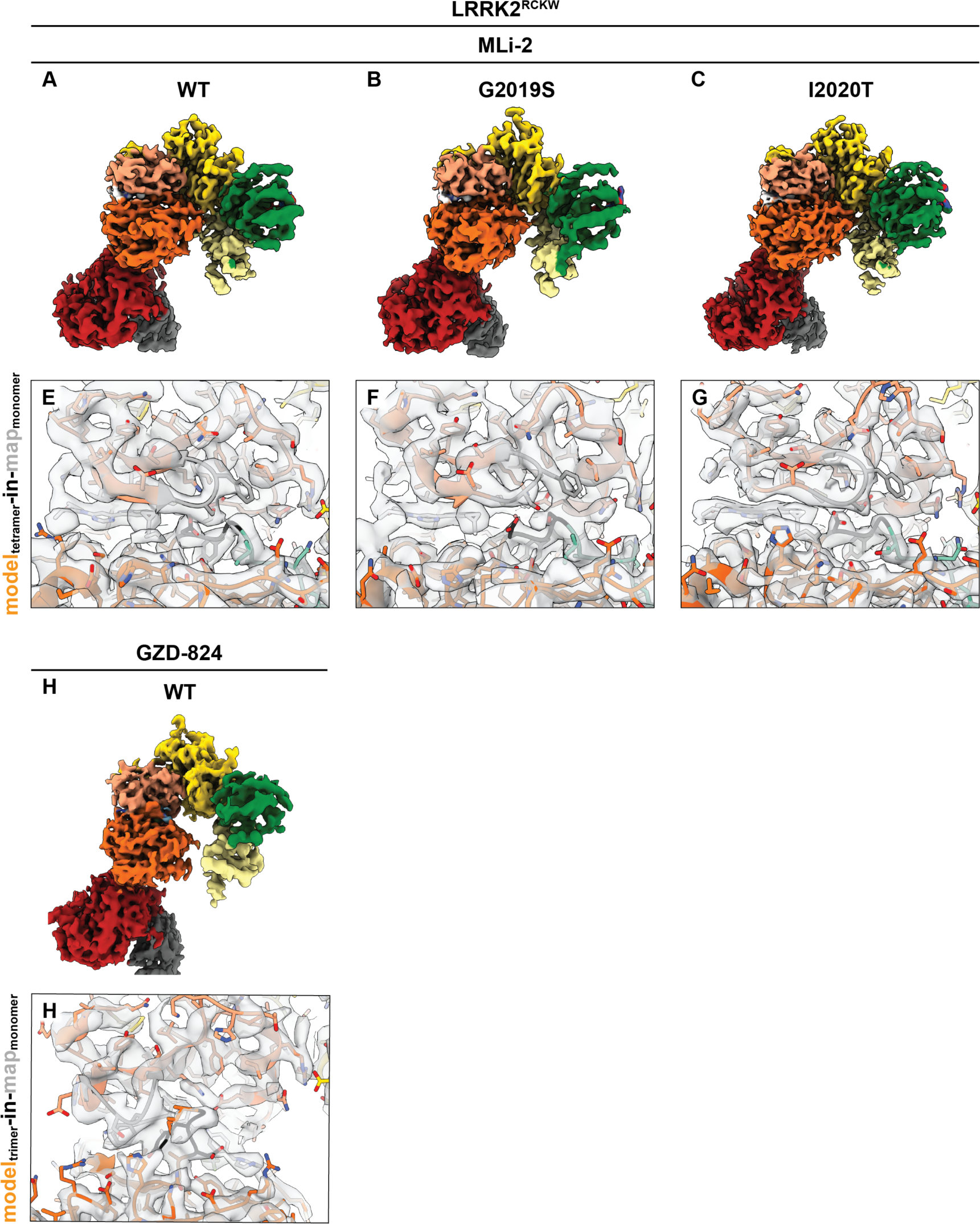
Cryo-EM maps of monomeric LRRK2^RCKW^’s account for models built into higher-resolution trimeric and tetrameric maps. **(A-D)** Cryo-EM maps of the monomeric form of LRRK2^RCKW^(WT):MLi-2 (A), LRRK2^RCKW^(G2019S):MLi-2 (B), LRRK2^RCKW^(I2020T):MLi-2 (C), and LRRK2^RCKW^(WT):GZD-824 (D). Maps are colored according to the domain color scheme shown in **Figure 1A**. The DARPin E11 is shown in dark grey. **(E-H)** The models built using the higher-resolution tetrameric (E-G) and trimeric (H) cryo-EM maps are fitted into the monomeric maps (see Methods for details on the cryo-EM data processing) to highlight that the kinase features discussed in the text (G-loop, DYG motif, αC helix, K1906-E1920 pair, and activation loop) are not dependent on the oligomeric state of LRRK2^RCKW^. The panels show the fit in and around the kinase’s active site.

**Figure S5.**
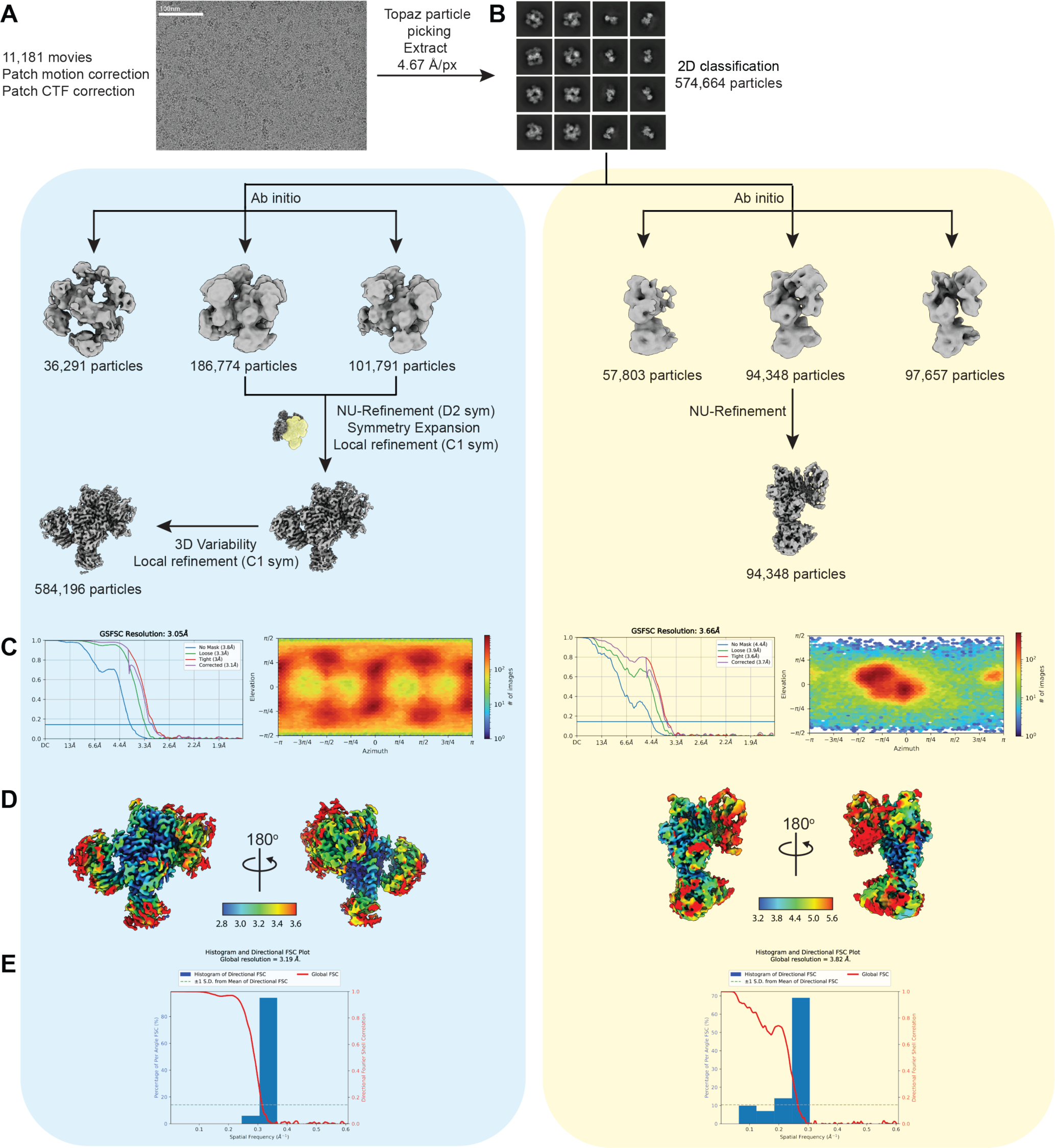
Cryo-EM workflow for LRRK2^RCKW^(WT):MLi-2. Representative micrograph **(A)**, 2D class averages and data processing strategy **(B)**, FSC plots and Euler angle distributions **(C)**, local resolution maps **(D)**, and 3D FSC plots **(E)** for LRRK2^RCKW^(WT):MLi-2 tetramer (blue panel) and monomer (yellow panel).

**Figure S6.**
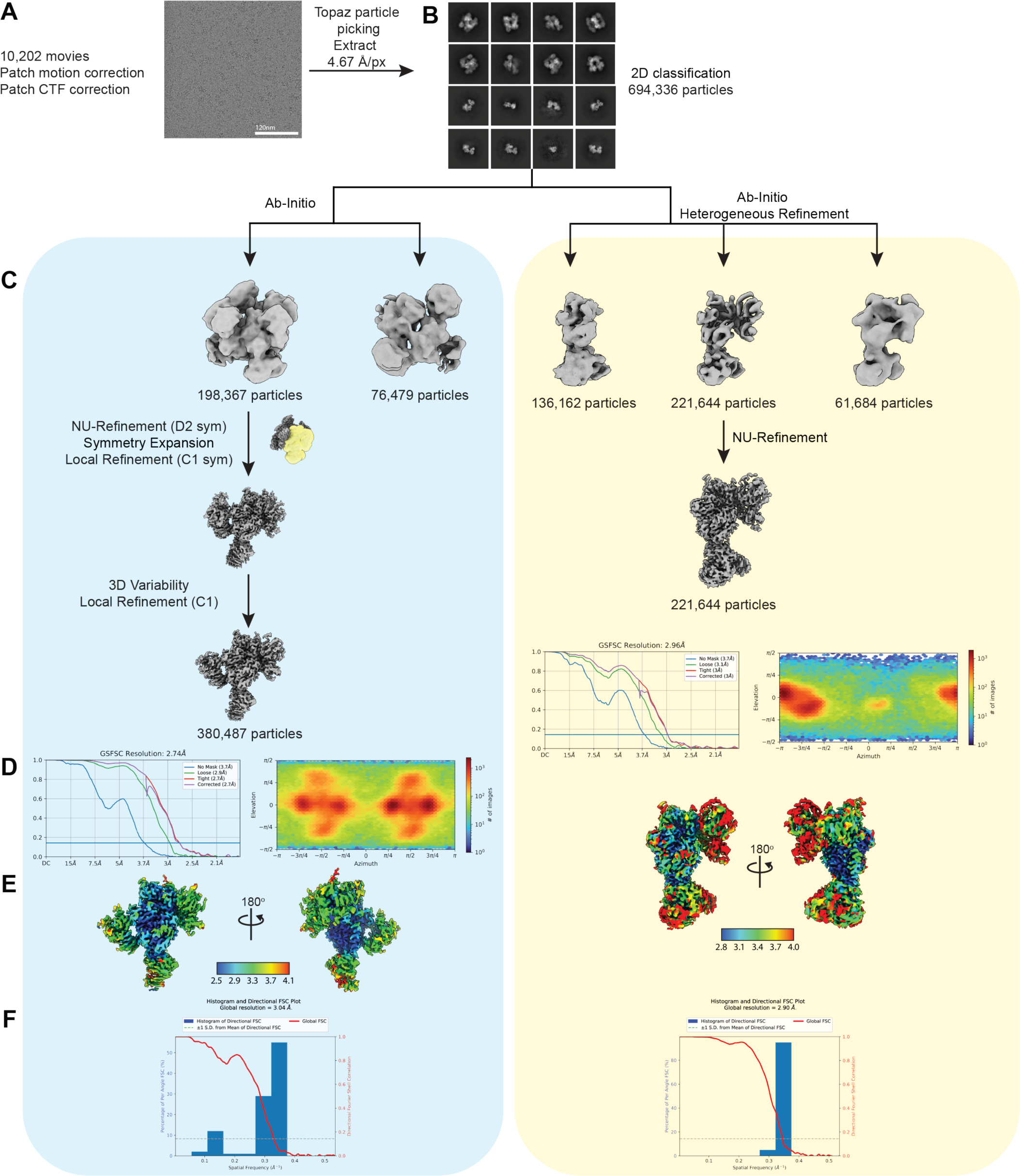
Cryo-EM workflow for LRRK2^RCKW^(I2020T):MLi-2. Representative micrograph **(A)**, 2D class averages **(B)**, data processing strategy **(C)**, FSC plots and Euler angle distributions **(D)**, local resolution maps **(E)**, and 3D FSC plot **(F)** for LRRK2^RCKW^(I2020T):MLi-2 tetramer (blue panel) and monomer (yellow panel).

**Figure S7.**
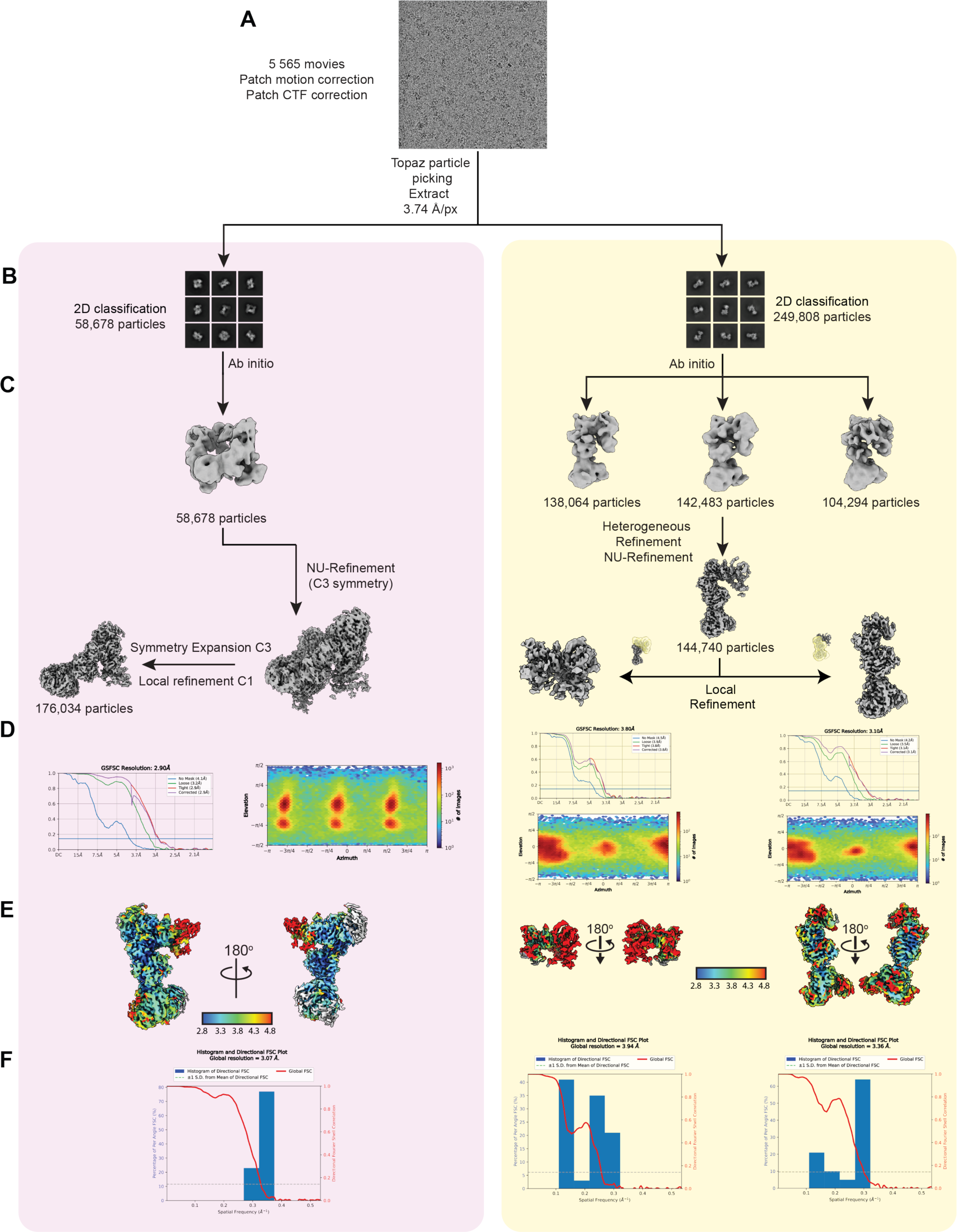
Cryo-EM workflow for LRRK2^RCKW^(WT):GZD-824. Representative micrograph **(A)**, 2D class averages **(B)**, data processing strategy **(C)**, FSC plots and Euler angle distributions **(D)**, local resolution maps **(E)**, and 3D FSC plot **(F)** for LRRK2^RCKW^(WT):GZD-824 trimer (pink panel) and monomer (yellow panel).

**Figure S8.**
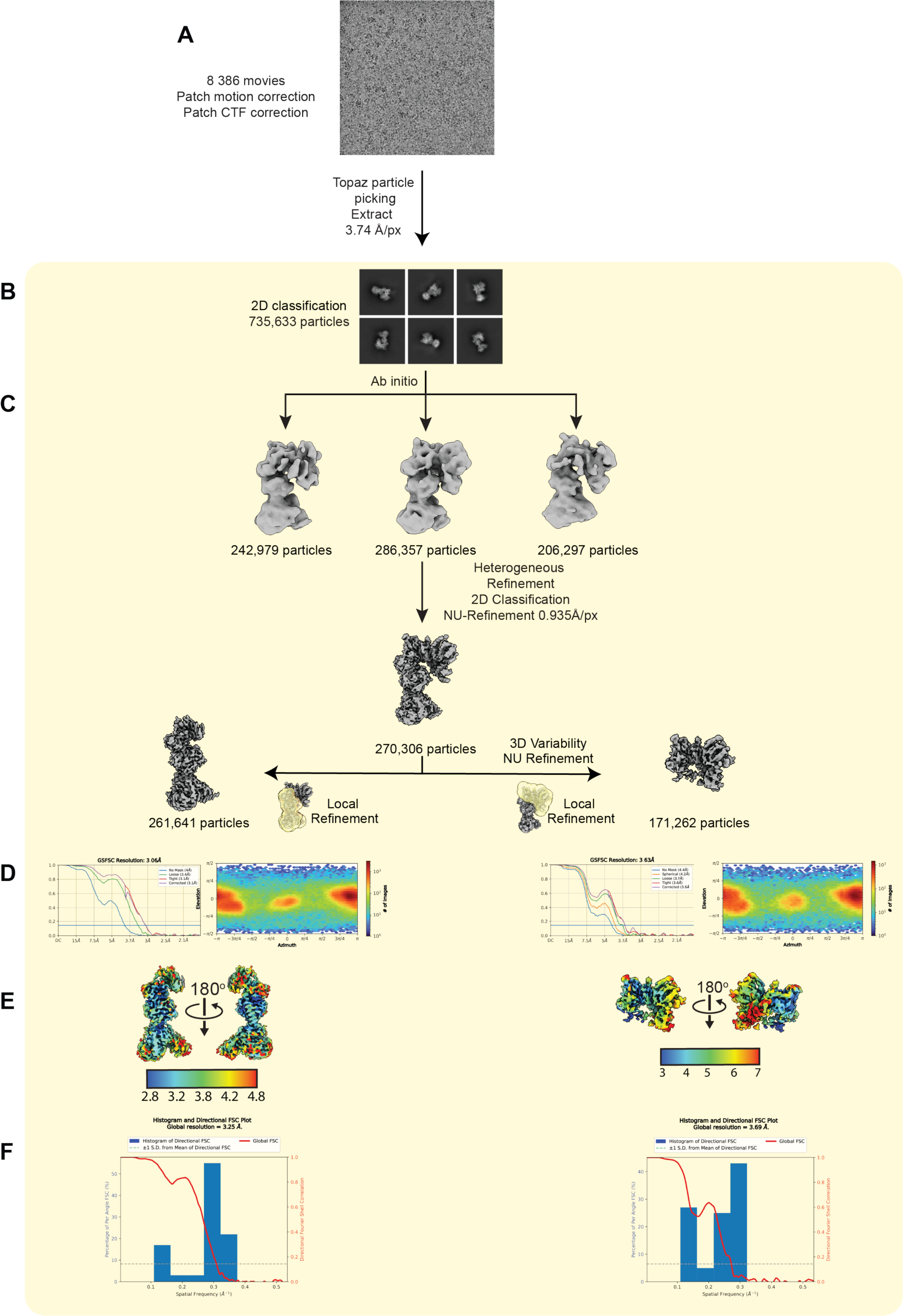
Cryo-EM workflow for LRRK2^RCKW^(I2020T):GZD-824. Representative micrograph **(A)**, 2D class averages **(B)**, data processing strategy **(C)**, FSC plots and Euler angle distributions **(D)**, local resolution maps **(E)**, and 3D FSC plot **(F)** for LRRK2^RCKW^(I2020T):GZD-824 monomer.

**Figure S9.**
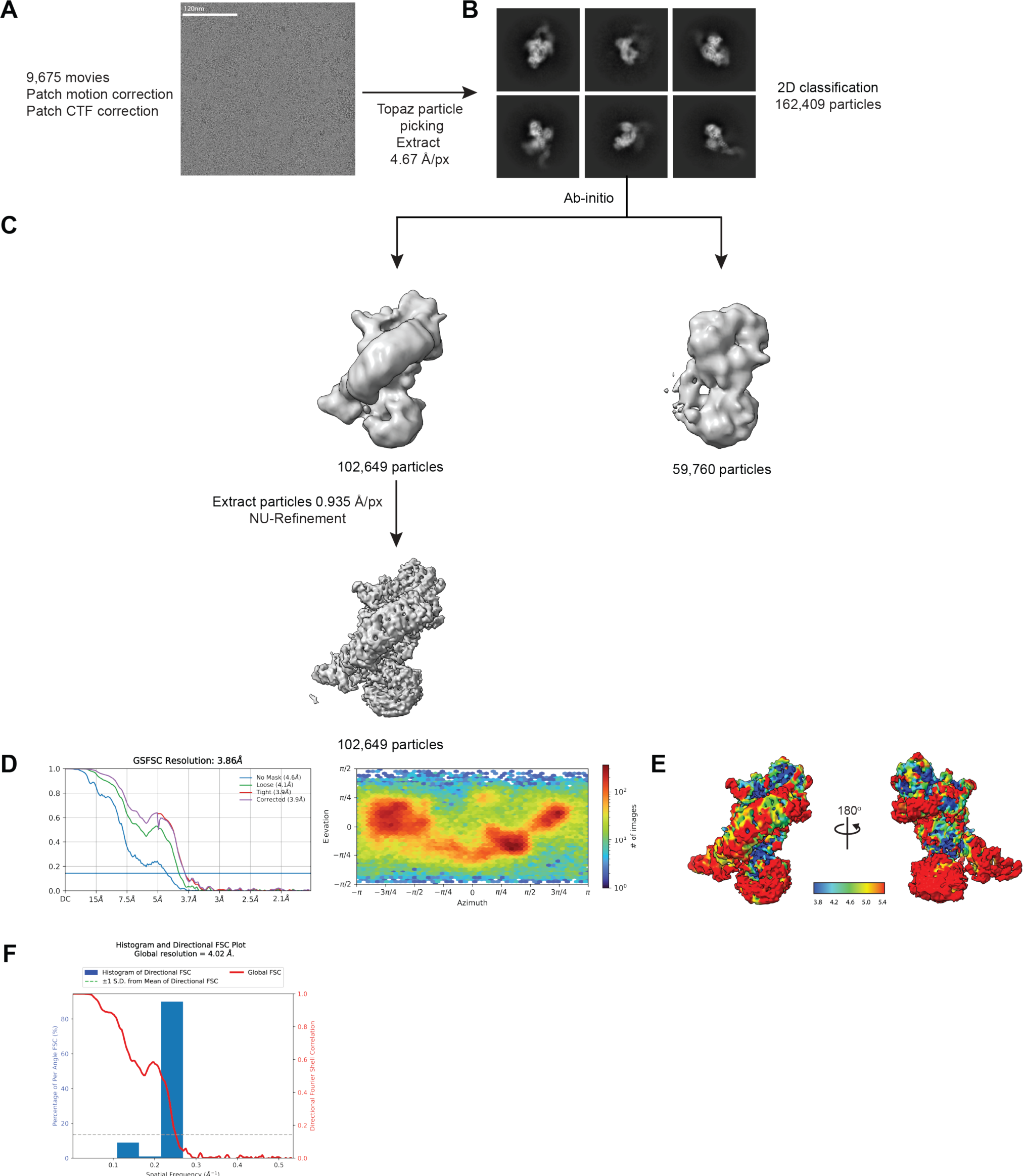
Cryo-EM workflow for LRRK2(I2020T):MLi-2. Representative micrograph **(A)**, 2D class averages **(B)**, data processing strategy **(C)**, FSC plots and Euler angle distributions **(D)**, local resolution maps **(E)**, and 3D FSC plot **(F)** for LRRK2(I2020T):MLi-2.

**Figure S10.**
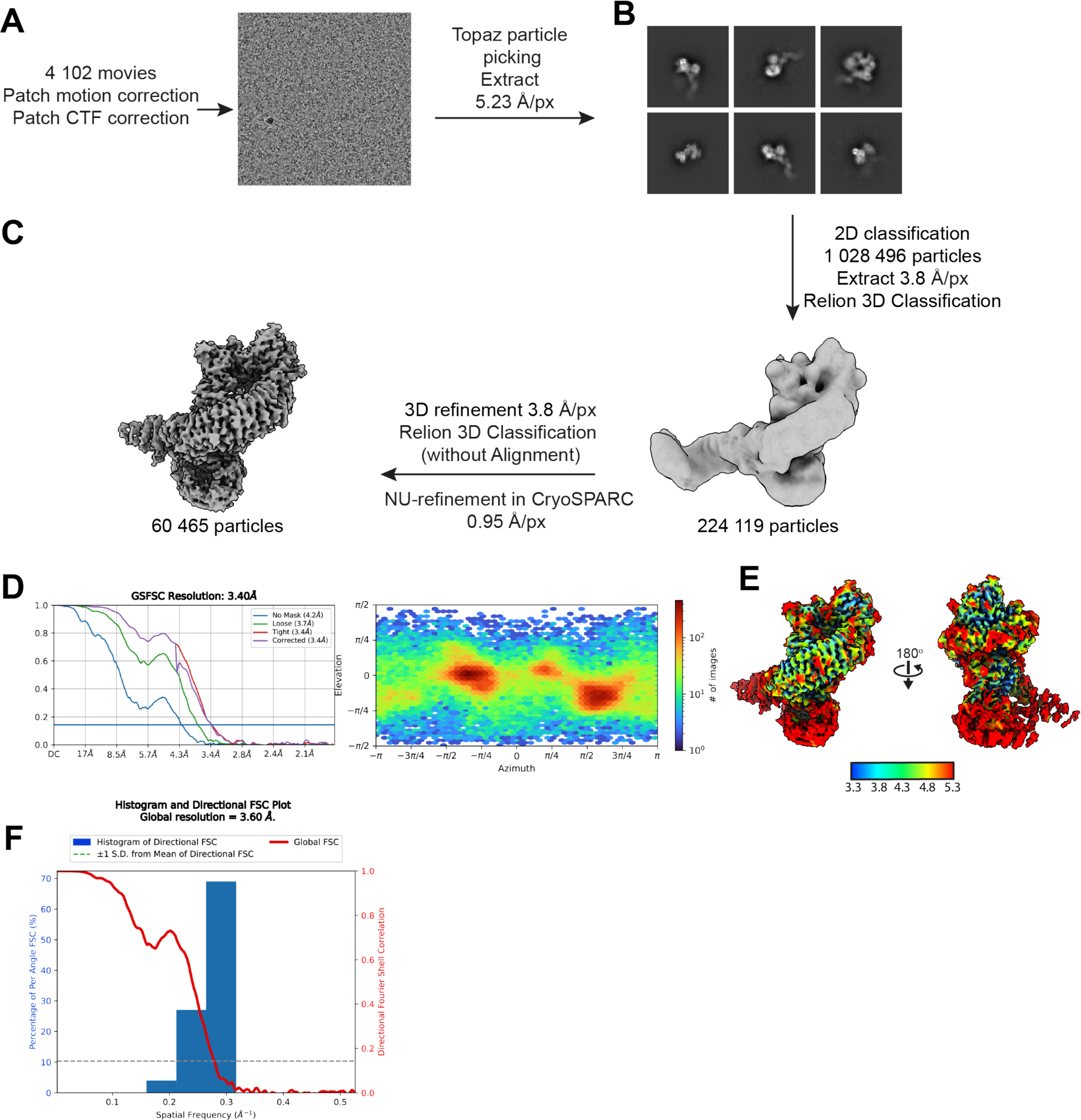
Cryo-EM workflow for LRRK2(I2020T):GZD-824. Representative micrograph **(A)**, 2D class averages **(B)**, data processing strategy **(C)**, FSC plots **(D)**, local resolution maps **(E)**, and 3D FSC plot **(F)** for LRRK2(I2020T):GZD-824.

**Movie S1. Structures of LRRK2 bound to type-1 and type-2 inhibitors**

The movie highlights the inhibitor poses and the major features of the LRRK2 kinase surrounding the type-1 and type-2 inhibitors in our structures of LRRK2^RCKW^(G2019S):MLi-2 and LRRK2^RCKW^(G2019S):GZD-824.

**Table S1.** Cryo-EM data collection, refinement, and validation statistics.

|  | LRRK2<sup>RCKW</sup>(WT):<br>MLi-2:E11 DARPin<br>(tetramer)<br>(EMDB-41709)<br>(PDB 8TXZ ) | LRRK2<sup>RCKW</sup>(G2019S):<br>MLi-2:E11 DARPin<br>(tetramer)<br>(EMDB-41754)<br>(PDB 8TZC) | LRRK2<sup>RCKW</sup>(I2020T):<br>MLi-2:E11 DARPin<br>(tetramer)<br>(EMDB-41758)<br>(PDB 8TZG) |
| --- | --- | --- | --- |
| **Data collection and processing** |  |  |  |
| Magnification | 105000 | 130000 | 130000 |
| Voltage (kV) | 300 | 300 | 300 |
| Electron exposure (e<sup>-</sup>/Å<sup>2</sup>) | 57 | 55 | 55 |
| Defocus range (μm) | -1.0 to -3.0 | -1.0 to -3.0 | -1.0 to -3.0 |
| Pixel size (Å) | 0.822 | 0.935 | 0.935 |
| Symmetry imposed | D2 | D2 | D2 |
| Initial particle images (no.) | 574 664 | 548 873 | 694 336 |
| Final particle images (no.) | 584 196 | 965 742 | 380 487 |
| Map resolution (Å) | 3.05 | 2.74 | 2.74 |
| FSC threshold | 0.143 | 0.143 | 0.143 |
| Map resolution range (Å) | 2.8 – 3.6 | 2.4 - 4.0 | 2.5-4.1 |
| **Refinement** |  |  |  |
| Initial model used (PDB code) | 6VP7 | 6VP7 | 6VP7 |
| Model resolution (Å) | 3.05 | 2.74 | 2.74 |
| FSC threshold | 0.143 | 0.143 | 0.143 |
| Model composition |  |  |  |
| Non-hydrogen atoms | 6261 | 7623 | 7095 |
| Protein residues | 814 | 998 | 907 |
| Nucleotide | 1 | 1 | 1 |
| Ligand | 1 | 1 | 1 |
| Water | 1 | 1 | 1 |
| *B* factors (Å<sup>2</sup>) |  |  |  |
| Protein | 42.37 | 93.28 | 47.54 |
| Nucleotide | 212.72 | 132.81 | 121.15 |
| Ligand | 23.59 | 78.41 | 34.58 |
| Water | 12.48 | 70.18 | 25.11 |
| R.m.s. deviations |  |  |  |
| Bond lengths (Å) | 0.019 | 0.008 | 0.003 |
| Bond angles (°) | 1.312 | 1.034 | 0.701 |
| Validation |  |  |  |
| MolProbity score | 2.30 | 2.21 | 1.83 |
| Clashscore | 8.85 | 7.97 | 8.06 |
| Ramachandran plot |  |  |  |
| Favored (%) | 93.92 | 93.74 | 94.31 |
| Allowed (%) | 5.96 | 6.26 | 5.58 |
| Disallowed (%) | 0.12 | 0.00 | 0.12 |

|  | LRRK2<sup>RCKW</sup>(G2019S):<br>GZD-824:E11<br>DARPin<sup>a</sup><br>(monomer)<br>Composite map<br>(EMDB-41728)<br>(PDB 8TYQ) | LRRK2<sup>RCKW</sup>(G2019S):<br>GZD-824:E11<br>DARPin<sup>a</sup><br>(monomer)<br>Consensus refine-<br>ment<br>(EMDB-41798) | LRRK2<sup>RCKW</sup>(G2019S):<br>GZD-824:E11<br>DARPin<sup>a</sup><br>(monomer)<br>Focused refinement<br>Kinase-WD40<br>(EMDB-41799) | LRRK2<sup>RCKW</sup>(G2019S):<br>GZD-824<br>E11<br>DARPin<sup>a</sup><br>(monomer)<br>Focused refinement<br>ROC-COR<br>(EMDB-41797) |
| --- | --- | --- | --- | --- |
| **Data collection and processing** |  |  |  |  |
| Magnification | 130000 | 130000 | 130000 | 130000 |
| Voltage (kV) | 300 | 300 | 300 | 300 |
| Electron exposure<br>(e-/Å<sup>2</sup>) | 55 | 55 | 55 | 55 |
| Defocus range (μm) | -1.0 to -3.0 | -1.0 to -3.0 | -1.0 to -3.0 | -1.0 to -3.0 |
| Pixel size (Å) | 0.935 | 0.935 | 0.935 | 0.935 |
| Symmetry imposed | C1 | C1 | C1 | C1 |
| Initial particle im-<br>ages (no.) | 759,293 | 759,293 | 759,293 | 759,293 |
| Final particle im-<br>ages (no.) | 261,641/169,878 | 261,641 | 261,641 | 169,878 |
| Map resolution (Å) | -- | 3.10 | 2.99 | 3.50 |
| FSC threshold | -- | 0.143 | 0.143 | 0.143 |
| Map resolution<br>range (Å) | 2.7-7 | 2.7-6 | 2.7-4.2 | 3-7 |
| **Refinement** |  |  |  |  |
| Initial model used<br>(PDB code) | 6VP7 | -- | -- | -- |
| Model resolution<br>(Å) | -- | -- | -- | -- |
| FSC threshold | -- | -- | -- | -- |
| **Model composition** |  |  |  |  |
| Non-hydrogen<br>atoms | 8523<br>1068 | --<br>-- | --<br>-- | --<br>-- |
| Protein residues | 0 | -- | -- | -- |
| Nucleotide | 1 | -- | -- | -- |
| Ligands |  |  |  |  |
| *B* factors (Å<sup>2</sup>) |  |  |  |  |
| Protein | 171.59 | -- | -- | -- |
| Nucleotide | -- | -- | -- | -- |
| Ligand | 48.03 | -- | -- | -- |
| **R.m.s. deviations** |  |  |  |  |
| Bond lengths (Å) | 0.007 | -- | -- | -- |
| Bond angles (°) | 0.904 | -- | -- | -- |
| **Validation** |  |  |  |  |
| MolProbity score | 1.72 | -- | -- | -- |
| Clashscore | 10.14 | -- | -- | -- |
| **Ramachandran plot** |  |  |  |  |
|  | 96.86 | -- | -- | -- |
| Favored (%) | 3.16 | -- | -- | -- |
| Allowed (%) | 0.00 | -- | -- | -- |
| Disallowed (%) |  |  |  |  |

<sup>a</sup> composite map

|  | LRRK2<sup>RCKW</sup>(I2020T):<br>GZD-824:E11 DARPin<br>(monomer)<br>Composite map<br>(EMDB- 41753)<br>(PDB 8TZB) | LRRK2<sup>RCKW</sup>(I2020T):<br>GZD-824:E11 DAR-Pin<sup>a</sup><br>(monomer)<br>Consensus refine-<br>ment<br>(EMDB- 41802) | LRRK2<sup>RCKW</sup>(I2020T):<br>GZD-824:E11<br>DARPin<sup>a</sup><br>(monomer)<br>Focused refinement<br>Kinase-WD40<br>(EMDB-41794) | LRRK2<sup>RCKW</sup>(I2020T):<br>GZD-824:E11<br>DARPin<sup>a</sup><br>(monomer)<br>Focused refinement<br>ROC-COR<br>(EMDB-41795) |
| --- | --- | --- | --- | --- |
| **Data collection and processing** |  |  |  |  |
| Magnification | 130000 | 130000 | 130000 | 130000 |
| Voltage (kV) | 300 | 300 | 300 | 300 |
| Electron exposure<br>(e-/Å<sup>2</sup>) | 55 | 55 | 55 | 55 |
| Defocus range (μm) | -1.0 to -3.0 | -1.0 to -3.0 | -1.0 to -3.0 | -1.0 to -3.0 |
| Pixel size (Å) | 0.935 | 0.935 | 0.935 | 0.935 |
| Symmetry imposed | C1 | C1 | C1 | C1 |
| Initial particle im-<br>ages (no.) | 735,633 | 735,633 | 735,633 | 735,633 |
| Final particle im-<br>ages (no.) | 261,641/171,262 | 270,306 | 261,641 | 171,262 |
| Map resolution (Å) | -- | 3.22 | 3.06 | 3.63 |
| FSC threshold | -- | 0.143 | 0.143 | 0.143 |
| Map resolution<br>range (Å) | -- | 2.8-7 | 2.8-4.4 | 3-7 |
| **Refinement** |  |  |  |  |
| Initial model used<br>(PDB code) | 6VP7 | -- | -- | -- |
| Model resolution<br>(Å) | -- | -- | -- | -- |
| FSC threshold | -- | -- | -- | -- |
| Model composition |  |  |  |  |
| Non-hydrogen<br>atoms | 7499<br>1012 | -- | -- | -- |
| Protein residues | 0 | -- | -- | -- |
| Nucleotide | 1 | -- | -- | -- |
| Ligands | -- | -- | -- | -- |
| *B* factors (Å<sup>2</sup>) |  |  |  |  |
| Protein | 145.4 | -- | -- | -- |
| Nucleotide | -- | -- | -- | -- |
| Ligand | 33.66 | -- | -- | -- |
| R.m.s. deviations |  |  |  |  |
| Bond lengths (Å) | 0.004 | -- | -- | -- |
| Bond angles (°) | 0.594 | -- | -- | -- |
| Validation |  |  |  |  |
| MolProbity score | 1.94 | -- | -- | -- |
| Clashscore | 10.46 | -- | -- | -- |
| Ramachandran |  |  |  |  |
| plot | 93.94 | -- | -- | -- |
| Favored (%) | 5.96 | -- | -- | -- |
| Allowed (%) | 0.11 | -- | -- | -- |
| Disallowed (%) | -- | -- | -- | -- |

<sup>a</sup> composite map

|  | FL(I2020T):<br>MLi-2:E11 DARPin<br>(monomer)<br>(EMDB-41759)<br>(PDB 8TZH) | FL(I2020T):<br>GZD-824: E11 DARPin<br>(monomer)<br>(EMDB-41757)<br>(PDB 8TZF) | LRRK2<sup>RCKW</sup>(WT):<br>GZD-824:E11 Dupin<br>(trimer)<br>(EMDB-41756)<br>(PDB 8TZE) |
| --- | --- | --- | --- |
| **Data collection and processing** |  |  |  |
| Magnification | 130000 | 150000 | 130000 |
| Voltage (kV) | 300 | 200 | 300 |
| Electron exposure (e-/Å<sup>2</sup>) | 50 | 55 | 55 |
| Defocus range (μm) | -1.0 to -3.0 | -1.0 to -3.0 | -1.0 to -3.0 |
| Pixel size (Å) | 0.935 | 0.95 | 0.935 |
| Symmetry imposed | C1 | C1 | C3 |
| Initial particle images (no.) | 162,409 | 224,119 | 58,678 |
| Final particle images (no.) | 102,649 | 60,465 | 176,034 |
| Map resolution (Å) | 3.9 | 3.4 | 2.90 |
| FSC threshold | 0.143 | 0.143 | 0.143 |
| Map resolution range (Å) | 3.8 -5.0 | 3.3-5.3 | 2.8-4.5 |
| **Refinement** |  |  |  |
| Initial model used (PDB code) | 7LHW | 7LHW | 6VP7 |
| Model resolution (Å) | 3.9 | 3.4 | 2.90 |
| FSC threshold | 0.143 | 0.143 | 0.143 |
| Model composition |  |  |  |
| Non-hydrogen atoms | 11,593 | 12,156 | 6251 |
| Protein residues | 1,606 | 1,671 | 787 |
| Nucleotide | 1 | 1 | 0 |
| Ligands | 1 | 1 | 1 |
| *B* factors (Å<sup>2</sup>) |  |  |  |
| Protein | 96.31 | 174.38 | 121.98 |
| Nucleotide | 102.80 | 87.23 | -- |
| Ligand | 82.68 | 109 | 45.07 |
| R.m.s. deviations |  |  |  |
| Bond lengths (Å) | 0.003 | 0.003 | 0.007 |
| Bond angles (°) | 0.791 | 0.557 | 1.096 |
| Validation |  |  |  |
| MolProbity score | 2.18 | 1.99 | 2.42 |
| Clashscore | 15.15 | 9.96 | 9.49 |
| Ramachandran plot |  |  |  |
| Favored (%) | 91.79 | 92.4 | 93.54 |
| Allowed (%) | 8.21 | 7.60 | 6.32 |
| Disallowed (%) | 0.00 | 0.00 | 0.13 |

